# Independent Activity Subspaces for Working Memory and Motor Preparation in the Lateral Prefrontal Cortex

**DOI:** 10.1101/756072

**Authors:** Cheng Tang, Roger Herikstad, Aishwarya Parthasarathy, Camilo Libedinsky, Shih-Cheng Yen

**Author notes:** Camilo Libedinsky and Shih-Cheng Yen contributed equally to this work.

## Abstract

The lateral prefrontal cortex is involved in the integration of multiple types of information, including working memory and motor preparation. However, it is not known how downstream regions can extract one type of information without interference from the others present in the network. Here we show that the lateral prefrontal cortex contains two independent low-dimensional subspaces: one that encodes working memory information, and another that encodes motor preparation information. These subspaces capture all the information about the target in the delay periods, and the information in both subspaces is reduced in error trials. A single population of neurons with mixed selectivity forms both subspaces, but the information is kept largely independent from each other. A bump attractor model with divisive normalization replicates the properties of the neural data. These results have implications for the neural mechanisms of cognitive flexibility and capacity limitations.

## Introduction

Complex flexible behaviors require the integration of multiple types of information, including information about sensory properties, task rules, items held in memory, items being attended, actions being planned, and rewards being expected, among others. A large proportion of neurons in the lateral prefrontal cortex (LPFC) encode a mixture of two or more of these types of information^1–5^. This mixed selectivity endows the LPFC with a high-dimensional representational space^1^, but it also presents the challenge of understanding how downstream regions that receive mixed-selective input from the LPFC can read out meaningful information. One possible solution would be to have multiple low-dimensional information subspaces, embedded within the high-dimensional state space of LPFC, which could enable the independent readout of different types of information with minimal interference from changes of information in other subspaces^6–10^. Information subspaces have been identified in the medial frontal cortex^11^, lateral prefrontal cortex^7^ and early visual areas^8^. However, no studies to date have explicitly tested whether two independent information subspaces can coexist within a single biological neural network. Here, we demonstrate the existence of two independent information subspaces in the LPFC network: one that encoded spatial working memory information, and another one that encoded movement preparation information. Both exhibited behavioral relevance with significantly decreased information in error trials only in the subspace, and not in the null space. Interestingly, we found a reduction in information in the memory subspace when information in the movement preparation subspace emerged. At the same time, we found that the average firing rate of the neurons across the population remained unchanged. This suggested that a normalization mechanism could have been acting on the population activity^12,13^. We subsequently found that a bump attractor model^14^ with divisive normalization allowed us to replicate the observed neurophysiological properties. We believe these results provide insights into the neural mechanisms of cognitive flexibility and cognitive capacity.

## Results

We measured LPFC activity while monkeys performed a delayed saccade task with an intervening distractor. Briefly, the monkeys had to remember the location (out of seven possibilities) of a briefly presented visual target for 2.3 seconds. One second after the target disappeared, a distractor was presented briefly in a different location. At the end of the 2.3 seconds, the monkeys reported the location of the remembered target using an eye movement (Fig. 1a). We measured the activity of 226 single units from the LPFC of both monkeys while they performed the task. We previously reported that the presentation of the distractor led to code-morphing in the LPFC, such that a decoder trained in the delay period that preceded the distractor (Delay 1) could not be used to decode memory information during the delay period that followed the distractor (Delay 2), and vice versa^2^ (Fig. 1b). In other words, there were two stable response periods, one in Delay 1 and one in Delay 2, but they did not generalize to each other.

**Figure 1.**
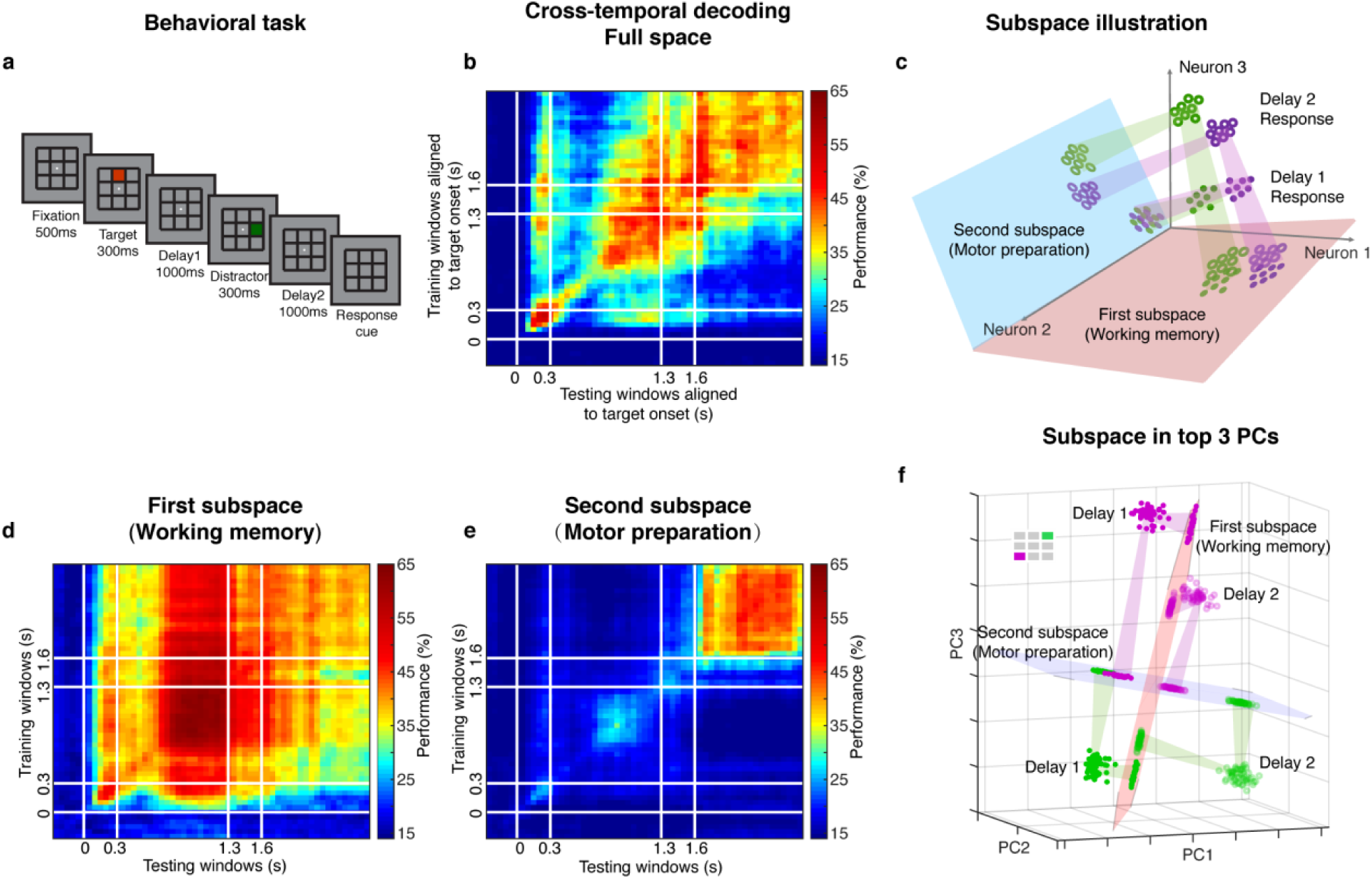
Experimental design, code morphing, and two independent subspaces. **a**, Behavioral task: Each trial began when the animal fixated on a fixation spot in the center of the screen. The animal was required to maintain fixation throughout the trial until the fixation spot disappeared. A target (red square) was presented for 300 ms followed by a 1000 ms delay period (Delay 1). A distractor (green square) was then presented for 300 ms in a random location that was different from the target location, and was followed by a second delay of 1000 ms (Delay 2). After Delay 2, the fixation spot disappeared, which was the Go cue for the animal to report, using an eye movement, the location of the target. **b**, Heat map showing the cross-temporal population-decoding performance in the LPFC. White lines indicate target presentation (0–0.3 s), distractor presentation (1.3–1.6 s), and cue onset (2.6 s). **c**, Schematic illustration of the projection of the full space activity into the *first* and *second subspaces*. Delay 1 activity (purple and green filled circles) projected into the *first subspace* would cluster according to target location (filled circles in the red subspace), and because this was a stable subspace, the Delay 2 activity for each target location (purple and green unfilled circles) would overlap with those for Delay 1 (open circles in the red subspace). In the *second subspace*, Delay 1 activity would not cluster according to location (filled circles in the blue subspace), and the clustering by location would emerge only from the Delay 2 activity (open circles in the blue subspace) after the emergence of the new information. **d**, Cross-temporal decoding performance after projecting full space activity into the memory subspace. **e**, Cross-temporal decoding performance after projecting full space activity into the preparation subspace. **f**, Projection of single trial activity for 2 target locations (actual locations shown in the upper left corner) onto the first 3 principal components. Delay 1 is depicted as closed circles, and Delay 2 as open circles. Re-projections into the *first subspace* (red plane) and *second subspace* (blue plane) are shown and guided by projection cones (green and purple cones connecting the PCA projections into the subspace re-projections).

### Two independent subspaces coexisted within the LPFC

The two different response patterns observed in Delay 1 and Delay 2 may imply that a downstream region would need to use two different decoders to read out the information, and would need to know which of them to use in the appropriate delay period to extract stable working memory information. Alternatively, the difference observed between Delay 1 and Delay 2 activity could be explained by an overlap of different types of information in independent subspaces, such that downstream regions can use different decoders to extract specific types of information. We have previously shown that a time-invariant (henceforth stable) working memory subspace can be identified in the LPFC^7^. However, significant information about the target was present outside of this space (null space decoding performance of 35.7 ± 1.7% in Delay 1, and 31.6 ± 1.5% in Delay 2)^7^ suggesting the existence of a non-trivial additional subspace that contains target information. In this paper, we investigated the possibility that: (1) the stable working memory subspace was one of two subspaces, in which target information emerged in Delay 1, and was maintained up until the end of Delay 2; and (2) information in the other subspace emerged only in Delay 2 after the presentation of the distractor, possibly due to the initiation of motor preparation after the last sensory cue that reliably predicted the timing of the Go cue (i.e. the offset of the distractor). The incorporation of the new information from the second subspace into the neuronal population, alongside the existing information from the first subspace, would have then resulted in code morphing in the full space (illustrated in Figure 1c). In order to assess this possibility, we used a novel method to identify the two subspaces. We first decorrelated the population activity in Delay 1 and Delay 2 by decomposing them into two components with the least mutual information, so that they could maximally capture different information (Supplementary Fig.1). Each component was a set of vectors that were the same dimensions as the population activity in the two delay periods (i.e. *226 x 7*, where 226 was the number of neurons, and 7 was the number of memory locations) in the full state space, and the basis vectors of the two components defined two subspaces (illustrated in Fig. 1c). Using this method, we found a pair of components that contained a minimum of 0.08 bits of mutual information (compared to 0.33 bits of mutual information in the activity of Delay 1 and Delay 2 prior to the decomposition). For the first component, the vector magnitude in Delay 2 was 65% of that in Delay 1. In contrast, the vector magnitude for the second component in Delay 1 was 12% of that in Delay 2 (the temporal dynamics of the population activity projected into these subspaces are shown in Supplementary Fig. 2). Cross-temporal decoding after projecting the neural activity into the *first subspace* showed that information emerged right after target presentation, and although the information was stronger in Delay 1 (60.5 ± 1.3%), it was present throughout the whole trial, even during the distractor period (Fig. 1d). This was qualitatively consistent with our hypothesis, aside from the decrease in information in Delay 2 (39.9 ± 1.1%). Cross-temporal decoding of neural activity projected into the *second subspace* showed that information emerged after distractor presentation (42.6 ± 1.1%), and was stable throughout Delay 2 (Fig. 1d). The subspaces had an effective dimensionality of 6 dimensions each, which accounted for more than 95% of the variance (Supplementary Fig. 2).

Figure 1f shows single trial projections of 2 different target locations (purple and green locations shown in the top-left corner) onto the top 3 principal components (PCs). These projections were then re-projected into the *first subspace* (red plane) and the *second subspace* (blue plane). Consistent with our hypothesis, Delay 1 and Delay 2 projections into the *first subspace* clustered according to target location, although they overlapped less than we expected (we will revisit this deviation from our expectation later on). However, the separation between the projected points was small enough that target location information could be decoded in both delays, regardless of whether the classifier was trained using Delay 1 or Delay 2 activity (Fig. 1d).

On the other hand, projections into the *second subspace* behaved differently, such that Delay 1 projections for multiple target locations overlapped, whereas Delay 2 projections remained separated. This explained why in the *second subspace*, target location information could not be decoded in Delay 1, but could be decoded in Delay 2 (Fig. 1e). Projections into the *first* and *second subspaces* for all target locations confirmed that these observations generalized to the rest of the locations (Supplementary Fig. 3, which also illustrates the reason for the difference in performance in the two off-diagonal quadrants in Fig. 1d).

### The two independent subspaces corresponded to working memory and motor preparation

Since the *first subspace* contained target information throughout the trial, and working memory of the target location was presumably required throughout the trial, we hypothesized that the *first subspace* corresponded to a working memory subspace. Using optimization methods, rather than decomposition as in the current study, we previously showed that the LPFC contained a working memory subspace that encoded stable working memory information^7^. In order to assess whether the *first subspace* corresponded to the working memory subspace previously described, we calculated the angle between these subspaces, as a measure of similarity (see Methods). We found that the *first subspace* was significantly closer than chance to the working memory subspace (*P* < 0.05, *g* = 2.62), while the *second subspace* was not (*P* > 0.28, *g* = 0.78, Supplementary Fig. 4). This result supported the interpretation that the *first subspace* corresponded to a working memory subspace. Thus, henceforth, we will refer to the *first subspace* as the “working memory subspace”.

Since the *second subspace* contained target information only after the distractor disappeared, and motor preparation presumably began after the last sensory cue that reliably predicted the timing of the Go cue (i.e. the offset of the distractor), we hypothesized that the *second subspace* corresponded to a motor preparation subspace. Activity between the Go cue and the saccade onset contained information about saccade execution (45% of LPFC neurons we recorded were selective in the period between the Go cue and saccade onset, assessed using a one-way ANOVA, *P* < 0.05). In order to test whether the *second subspace* corresponded to a motor preparation subspace, we compared the results of the original decomposition using Delay 1 and Delay 2 activities with a new decomposition using Delay 1 and pre-saccadic period activities (150 ms to 0 ms prior to saccade). If the *second subspace* corresponded to a motor preparation subspace, we should observe similarities between the second component in both decompositions. In the new decomposition, we obtained two components with relative vector magnitudes similar to those found in the first decomposition (For *Component 1’*, the vector magnitude in the pre-saccade period was 70% of that in Delay 1, while for *Component 2’*, the vector magnitude in Delay 1 was 0% of that in the pre-saccade period, Supplementary Fig. 5). We found that *Component 1* and *Component 1’* were significantly correlated (Fig. 2a left, Pearson correlation *r* > 0.99, *P* < 0.01). Importantly, *Component 2* and *Component 2’* were also significantly correlated (Fig. 2a right, Pearson correlation *r* = 0.62, *P* < 0.01). This result supported our hypothesis that the *second subspace* corresponded to a motor preparation subspace.

**Figure 2.**
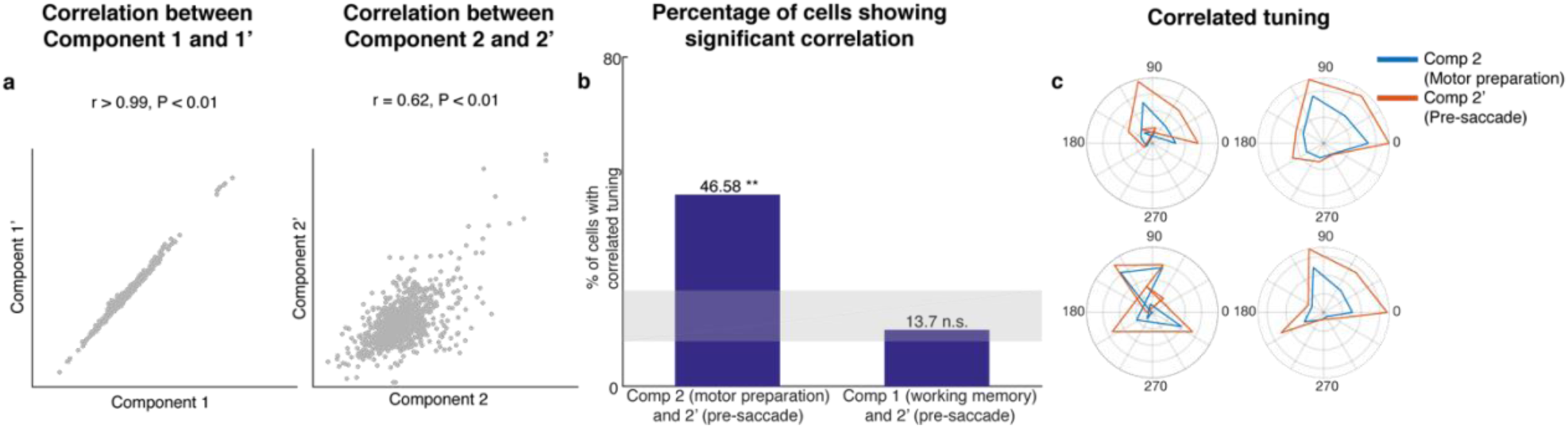
Preparatory and pre-saccadic activity. **a**, Correlation between components found in the in the decomposition of Delay1 / Delay 2 activities (Components 1 and 2) and the decomposition of Delay / pre-saccadic activities (Components 1’ and 2’). **b**, Left, the percentage of cells that exhibited significant correlation between *Components 2* and *2’*. Right, the percentage of cells that exhibited significantly correlation between *Components 1* and *2’*. The shaded area shows the 5th and 95th percentiles of the chance percentage obtained by shuffling the tuning across cells. **c**, Response tuning of *Component 2 and 2’* of four representative cells that showed significant correlation.

In an additional test of the hypothesis that the *second subspace* corresponded to a motor preparation subspace, we examined the relationship between *Component 2* and *Component 2’* at the level of single cells. First, we identified cells with spatial tuning in both Delay 2 and the pre-sacade period (73 cells, two one-way ANOVAs, both *P* < 0.05). Then, for each cell, we measured the correlation between the activity in *Component 2* and *Component 2’* across different target locations. We found that 47% of these neurons showed significant correlation (Pearson correlation, *P* < 0.05), which exceeded the number expected by chance (Fig. 2b, left bar, *P* < 0.001, *g* = 10.82). As a control, we carried out the same analysis between *Component 1* and *Component 2’*, and found no evidence of a higher number of correlated cells than expected by chance (Fig. 2b, right bar, *P* > 0.19, *g* = 1.51). Examples of neurons with significant correlation are shown in Figure 2c. This result provided additional support to our hypothesis that the *second subspace* corresponded to a motor preparation subspace. Thus, henceforth, we will refer to the *second subspace* as the “motor preparation subspace”.

### Responses of neurons with mixed working memory and motor preparation selectivity formed the two subspaces

The existence of two independent subspaces could be mediated by one of two possible mechanisms: (1) two independent subpopulations of neurons with exclusive working memory or motor preparation selectivity within the LPFC, or (2) the same population of LPFC neurons with mixed selectivity to both working memory and motor preparation. In order to distinguish between these two possible mechanisms, we projected the unit vector representing each neuron in the full space into the working memory and motor preparation subspaces, and quantified the magnitude of the two projections for each neuron (i.e. loading weight, Fig. 3a). An anticorrelation between the loading weights in each subspace would support the first mechanism, and a positive correlation would support the second mechanism. We found a significant positive correlation between the loading weights in each subspace (*r* = 0.68, *P* < 0.001), in support of the second mechanism. As a measure of the relative contribution to each subspace, we calculated the ratio between the loading weights for each cell, and analyzed their distribution (Fig. 3b). We found that only 14 (6%) of the neurons had “exclusive” loading for the working memory (red) or motor preparation (blue) subspaces. However, these cells were not necessary to identify the subspaces (Supplementary Fig. 6).

**Figure 3.**
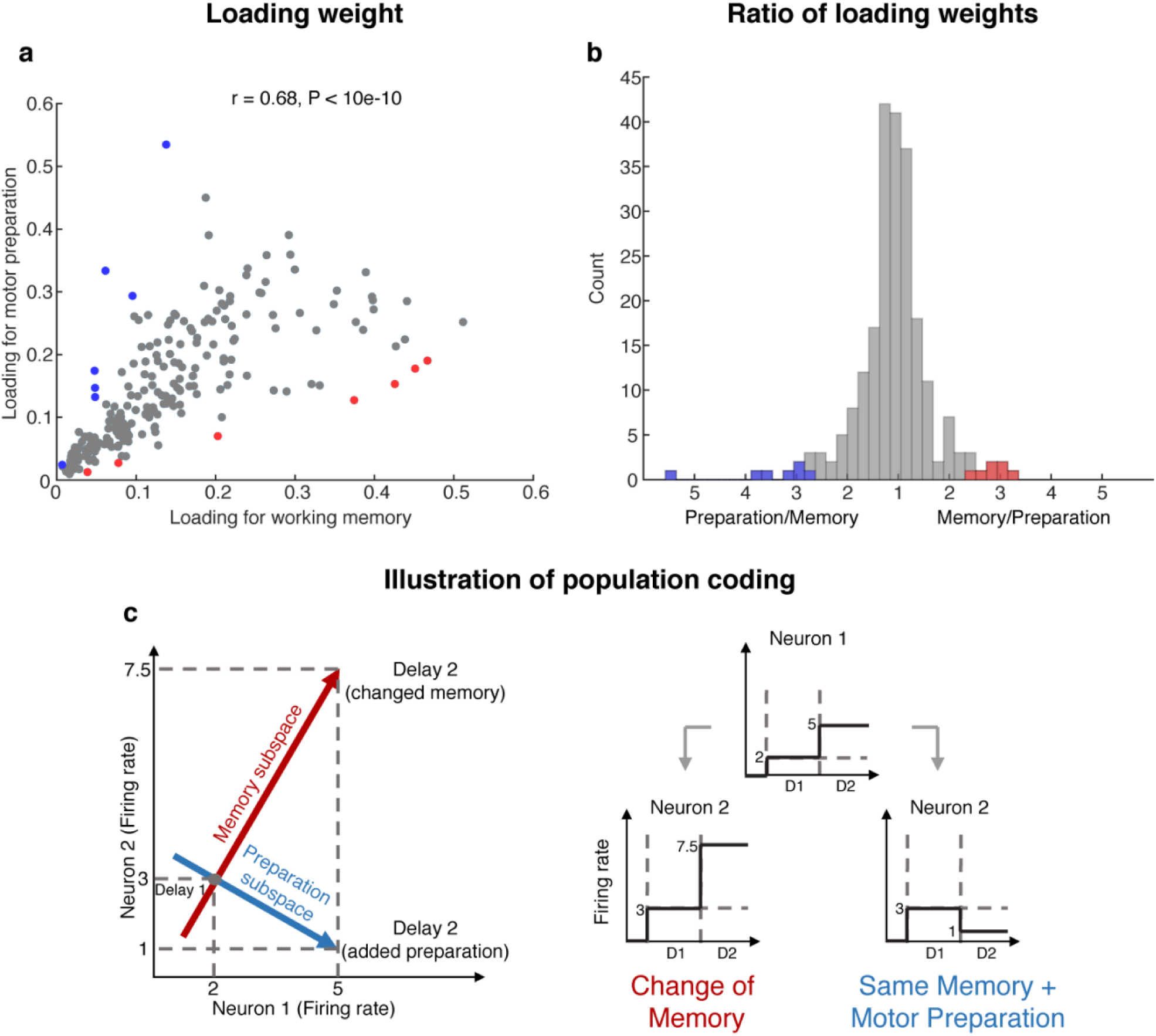
Loading weights for individual neurons. **a**, The loading weight of each neuron in the working memory subspace and the motor preparation subspace. **b**, Histogram of the ratio between the loading weights for each cell. For cells with larger loading for the working memory subspace, the values are plotted to the right of the plot, while for cells with larger loading for the motor preparation subspace, the values are plotted to the left of the plot. Red dots (in **a**) and bars (in **b**) represent cells with “exclusive” loading for the working memory subspace, while blue dots (in **a**) and bars (in **b**) represent cells with “exclusive” loading for the motor preparation subspace. These cells were identified as those with ratios that exceeded 2 standard deviations from the mean. **c**, Illustration of population coding. The conjunctive population code formed by both Neuron 1 and Neuron 2 unambiguously direct information in the memory subspace or preparation subspace in Delay 2, whereas it would have been ambiguous to look only at Neuron 1’s change of firing rate in Delay 2.

In order to understand how a single population of neurons with mixed selectivity could have contributed to two independent subspaces, we created a simple illustration (Fig. 3c). In isolation, the activity of Neuron 1 would be ambiguous, as an increase of activity in Delay 2 could be interpreted as a new memory at a different spatial location, or as the same memory as in Delay 1, but with overlapping motor preparation activity. In order to disambiguate the meaning of a change in the activity of one neuron, it would be necessary to interpret that change in the context of changes in the activity of the rest of the neuronal population (i.e. in this example, Neuron 2). In the illustration, a concurrent increase of activity in Neurons 1 and 2 signals a change in memory (i.e. the population activity in state space moved along the working memory subspace), whereas the same increase in Neuron 1, but with a concurrent decrease of activity in Neuron 2, signals that the memory has not changed, but that a motor preparation plan has emerged in Delay 2 (i.e. the population activity in state space moved along the motor preparation subspace). This concept can be extended to the 212 neurons with mixed selectivity to understand how coordinated activity between those neurons can unambiguously contribute information to the independent 6-dimensional working memory and motor preparation subspaces that we found in the LPFC.

### Information in one subspace led to a small amount of interference in information in the other subspace

Since one population of neurons with mixed selectivity contributed to both the working memory and motor preparation subspaces, it was possible that information in one of the subspaces interfered with information in the other, and vice versa. One way to assess the degree of interference between subspaces was to quantify the drop in decoding performance when activity related to both working memory and motor preparation overlapped (i.e. Delay 2). In order to quantify this interference, we compared the decoding performance of a classifier trained and tested on working memory activity projected into the working memory subspace (Fig. 4a: proj_MSub_(M)), with decoding performance of a classifier trained and tested on working memory and motor preparation activity (i.e. full space activity) projected into the working memory subspace (Fig. 4a: proj_MSub_(M+P)). A similar analysis was carried out in the motor preparation subspace (Fig. 4b: proj_PSub_(P) and proj_PSub_(M+P)). We found no evidence of a drop in performance between proj_MSub_(M) and proj_MSub_(M+P) (*P* > 0.85, *g* = 0.66), and between proj_PSub_(P) and proj_PSub_(M+P) (*P* > 0.15, *g* = 2.64), suggesting a lack of interference between these subspaces.

**Figure 4.**
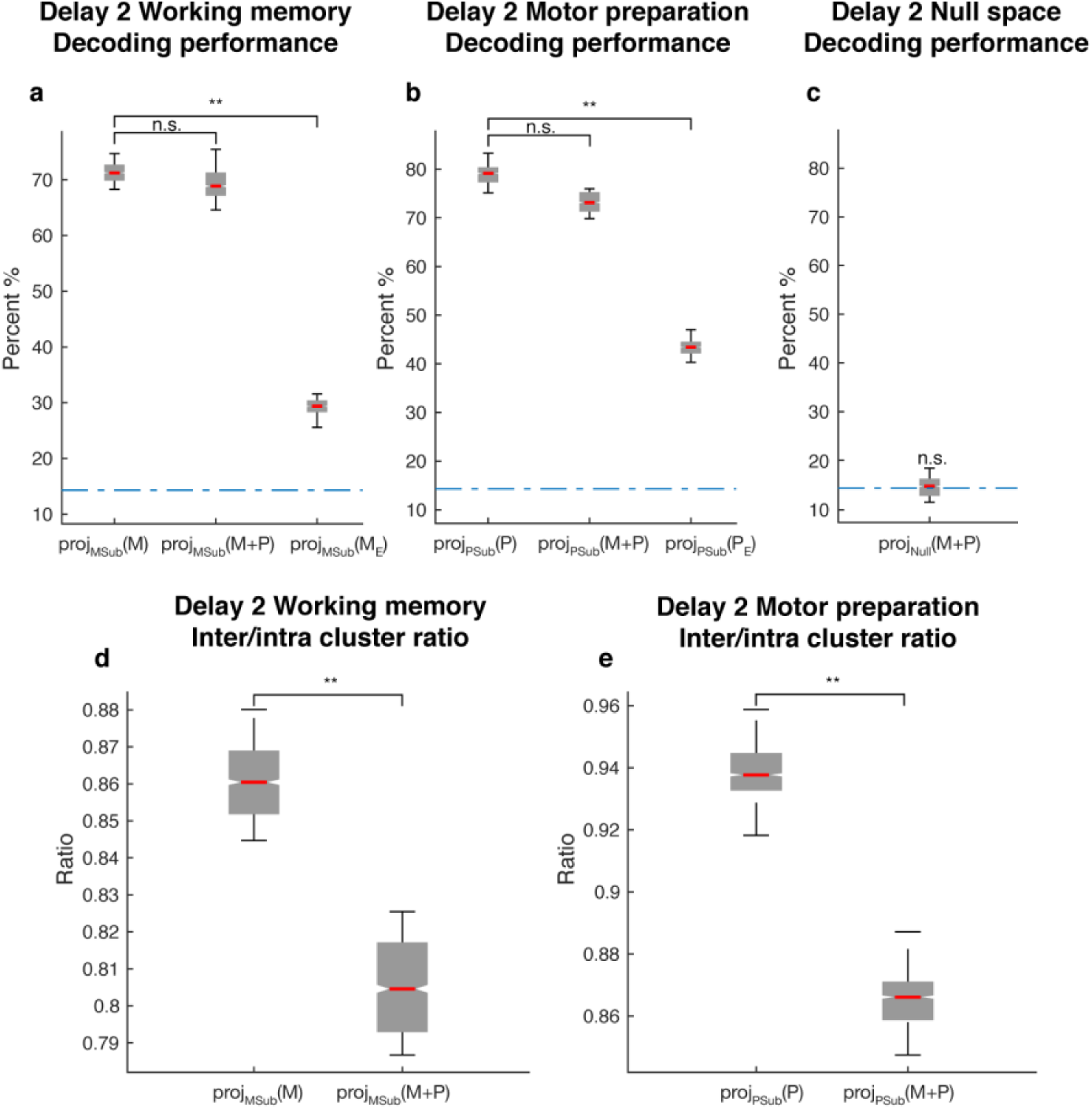
Comparisons between working memory and motor preparation subspaces. **a**, *M* stands for working memory activity; proj_MSub_(M), decoding of the working memory activity projected into the working memory subspace; proj_MSub_(M+P), decoding of the full space activity projected into the working memory subspace; proj_MSub_(M_E_), decoding of the working memory activity in error trials projected into the working memory subspace using a classifier built on working memory activity in correct trials projected into the working memory subspace; **b**, *P* stands for motor preparation activity. Same conventions as in **a**, but for motor preparation activity and the motor preparation subspace. We verified that the drop in performance in error trials was specific to the two subspaces, and not due to a non-specific increase in noise in the population (see Methods). **c**, proj_Null_(M+P), decoding of the full space activity projected into the null subspace. **d**, Inter-to-Intra cluster ratio of working memory activity projected into the working memory subspace (proj_MSub_(M)), and of full space activity projected into the working memory subspace (proj_MSub_(M+P)). **e**, Same conventions as in **d**, but for motor preparation activity and motor preparation subspace.

The lack of a difference in decoding performance did not rule out the possibility that there was a small interference that did not significantly affect the classifier’s performance, so we performed a more sensitive state-space analysis on Delay 2 activity to assess whether the working memory and motor preparation subspaces interfered with each other. We quantified interference by projecting the activity of single trials into the two subspaces, and calculated the average distance between clusters of points corresponding to different target locations (inter-cluster distance). The inter-cluster distance was then normalized by the average intra-cluster distance for all clusters, which was a measure of trial-by-trial variability in the population response. This inter-to-intra cluster distance ratio was compared between projections of working memory activity into the working memory subspace (Fig. 4d: proj_MSub_(M)), and projections of working memory and motor preparation activity into the working memory subspace (Fig. 4d: proj_MSub_(M+P)). A similar analysis was carried out in the motor preparation subspace (Fig. 4e). We found a small decrease (7.1%) of the inter-to-intra cluster distance when both working memory and motor preparation components were projected into the subspaces, suggesting the existence of a small, but significant interference between both subspaces (*P* < 0.001, *g* = 4.81 between proj_MSub_(M) and proj_MSub_(M+P), *P* < 0.001, *g* = 6.63 between proj_PSub_(P) and proj_PSub_(M+P)).

### The two subspaces accounted for all the target information in the full space

We identified 2 independent subspaces that accounted for working memory and motor preparation signals. However, it was possible that there were other subspaces that also contained target location information. If this was true, information about target location should be present in the subspace that complemented the 6-dimensional working memory and the 6-dimensional motor preparation subspaces (i.e. the null space that has 117 effective dimensions that explain 95% total variance). In order to assess this possibility, we trained and tested a decoder on activity projected into the null space. We found that the null space contained no information (Fig. 4c, proj_Null_(M+P), *P* > 0.59, *g* = 0.01 compared to chance), which meant that the two subspaces captured all the decodable information in the full space. This observation suggested that the 2 subspaces we identified were the only ones that contained target information in the LPFC in our task.

### Less information was found in error trials in both subspaces

In a subset of trials, the animals maintained fixation until the Go cue, but failed to report the correct target location with a saccade. These failures could be due to the animals reporting other locations, including the location of the distractor, or simply saccading to a completely different location, such as the edge of the monitor. Classifiers trained on working memory activity of correct trials projected into the working memory subspace, proj_MSub_(M), were tested on working memory activity of error trials, which was also projected into the working memory subspace, proj_MSub_(M_E_). Decoding performance was significantly reduced in error trials (Fig. 4a,) compared to correct trials (Fig. 4a, *P* < 0.001, *g* = 14.8 between proj_MSub_(M) and proj_MSub_(M_E_)), suggesting that failures in memory encoding occurred during error trials. A similar analysis on the motor preparation subspace yielded equivalent results (Fig. 4b, *P* < 0.001, *g* = 14.8 between proj_PSub_(P) and proj_PSub_(P_E_)), which were consistent with the fact that in error trials, saccades were made to different locations than in correct trials. These results suggested that the subspaces we found could have been used by the animals to perform the task.

### A bump attractor artificial neural network with divisive normalization recapitulated the properties of LPFC activity

An unexpected observation in our results was that decoding performance in the working memory subspace decreased in Delay 2 compared to Delay 1 (Fig. 1d). This decrease coincided with an increase of decoding performance in the motor preparation subspace (Fig. 1e). A state space analysis revealed that the decrease of working memory decoding performance was due to a decrease in the inter-to-intra cluster distance of working memory signals in Delay 2, and the increase of motor preparation decoding performance was due to an increase in the inter-to-intra cluster distance of motor preparation signals in Delay 2, (Supplementary Fig. 7). In addition, we noticed that the mean population firing rate did not change between Delay 1, Delay 2 and pre-target fixation periods (Supplementary Fig. 8). This observation was consistent with a population normalization mechanism to maintain the mean population firing rate at a constant level in the LPFC^12,13^. In order to assess whether a normalization mechanism was responsible for the decrease of working memory information in Delay 2, we built artificial neural network models with and without population normalization, and compared their behavior with the LPFC data.

Bump attractor models have been shown to replicate several properties of LPFC activity, including code-morphing in full space, the existence of a stable subspace with stable working memory information, and non-linear mixed selectivity of individual neurons^7,14,15^. Here, we created a model that incorporated subsets of neurons that represented information in the working memory and motor preparation subspaces (Fig. 5a), and looked at the effect of adding divisive normalization to keep the mean population firing rate constant. We constrained the model to utilize neurons with mixed selectivity (Fig. 3a) by matching the selectivity properties to those found in the LPFC data (Supplementary Fig. 9). We also used the same decorrelation method to identify working memory and motor preparation subspaces from the responses in the model (Supplementary Fig. 10). In both models with and without normalization, a stable working memory bump appeared in Delay 1 after a subset of neurons representing one of the working memory locations was activated (shown in the bottom left of Fig. 5b and 5d, where the cross-temporal decoding performance in *LP11* was 75.8 ± 6.9% and 77.0 ± 9.6% respectively). After a second subset of neurons representing motor preparation were activated in Delay 2, the model without normalization exhibited an elevated decoding performance in the full space (93.1 ± 3.5% in *LP22*, Fig. 5b) due to the correlated location between working memory and motor preparation, which increased the distances between the clusters in the state space (Fig. 5c). There was also no reduction of performance in the working memory subspace in Delay 2 (76.5 ± 8.3% in *LP11*, 77.8 ± 8.1% in *LP22*, *P* > 0.86, *g* = 0.28), and the mean population activity increased from Delay 1 (1.2 ± 0.04 spikes/s) to Delay 2 (1.4 ± 0.06 spikes/s, *P* < 0.05, *g* = 4.04). These three results were inconsistent with our observations from the neuronal data. On the other hand, in the model with normalization, we found that the decoding performance in the full space remained the same in both delay periods (*LP22* -*LP11* overlapped with 0, *P* > 0.78, *g* = 0.61, Fig. 5d), replicated the reduction in information in the working memory subspace in Delay 2 (82.6 ± 9.4% in *LP11*, 56.1 ± 5.4% in *LP22*, P < 0.05, g = 3.42, Fig. 5e), and did not observe any changes in the mean population firing rate (D1: 1.0 ± 0.04 spikes/s, D2: 1.0 ± 0.04 spikes/s, P > 0.9, g = 0.05). Finally, as expected, target information in the motor preparation subspace emerged in Delay 2 (Fig. 5f). A linear subspace model^16^ with divisive normalization was also able to replicate qualitatively all the important findings of the neural data (see Methods and Supplementary Fig. 11), indicating the conceptual convergence between the bump attractor model and the linear subspace model.

**Figure 5.**
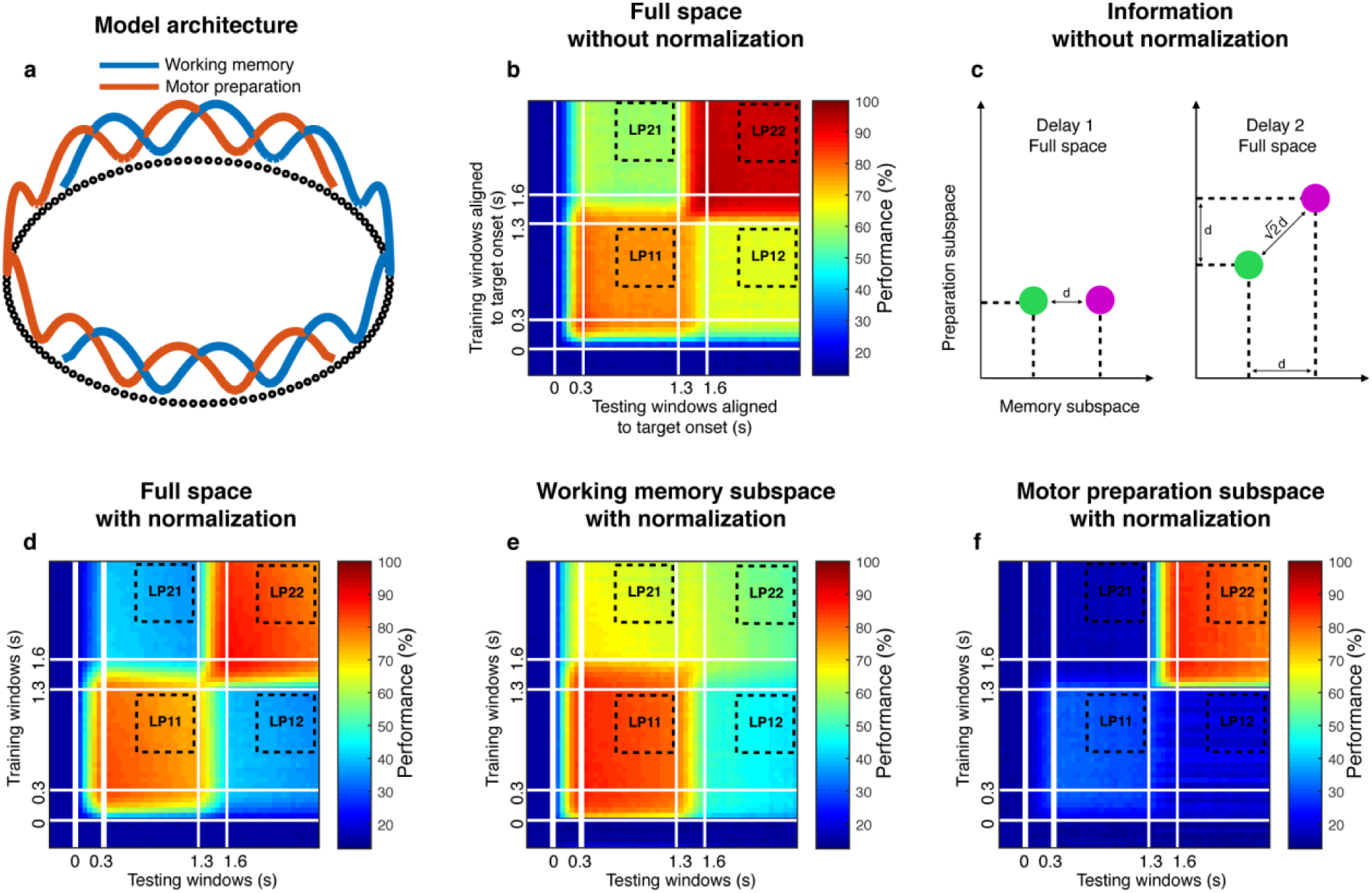
Neural network model with and without normalization. **a**, Bump attractor model (adapted from Wimmer et al., 2014^15^) with overlapping working memory and motor preparation populations. The full population (112 units) received two sets of inputs: 8 working memory and 8 motor preparation inputs. Each input activated sets of 10 adjacent units, and the working memory and motor preparation inputs overlapped by 43% (which matched the percentage of overlap in the LPFC data). **b**, Cross-temporal decoding of the model (without normalization) in the full space. **c**, Schematic illustrating the increase in information without normalization. Green and purple circles represent two different target clusters in the full space separated by an inter-cluster distance of *d*. If preparation information (in the form of inter-cluster distance of *d*) was added, this would result in the full space inter-cluster distance in Delay 2 increasing by a factor of 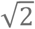. **d**, Cross-temporal decoding performance of the model (with normalization) in the full space. **e**, Cross-temporal decoding performance of the model (with normalization) in the working memory subspace. **f**, Cross-temporal decoding performance of the model (with normalization) in the motor preparation subspace.

## Discussion

Here we demonstrate that two independent subspaces coexist within the LPFC population. These subspaces contain largely independent information about target location, and appear to encode working memory and motor preparation information. We show that there is a small, but significant interference of information when both subspaces encode information simultaneously, and during error trials information in the working memory subspace is reduced. Assessment of response properties of individual neurons revealed that a single population of neurons with mixed selectivity generates both subspaces. Finally, we show that a bump attractor neural network model with divisive normalization can capture all these properties described. Overall, our results show that working memory and motor preparation subspaces coexist in a single neural network within the LPFC.

It is important to minimize interference between different types of information. For example, a visual area may read out working memory information^17,18^, while a premotor region may read out motor preparation information from the LPFC^17,19,20^. If large interferences existed between subspaces, the computations of downstream regions would be compromised. We found a small, but significant interference between the subspaces, such that some working memory information was reflected in the motor preparation subspace (and vice versa). It is not surprising that there is some degree of interference, since the method we used to decompose the signals did not impose a constraint to ensure maximal orthogonality between subspaces, and while the mutual information was low, it was not zero. To assess whether imposing orthogonality between subspaces was feasible, we attempted to decompose the signals by minimizing the inner product rather than the mutual information. However, this method did not provide a unique solution for the decomposition, and the best decoding performance and interference obtained by this method was not significantly different from that obtained by the method of minimum mutual information (Supplementary Fig. 12). We also considered alternative methods to decompose the signal, but these produced subspaces with larger interferences (Supplementary Figs. 13 and 14). The interference we found suggests that under conditions that stress the working memory and motor preparation systems (such as a task that requires the concurrent memorization of 4 targets and preparation of 4 movements) a predictable bias should be observable for both the recalled target locations and eventual movements. This prediction remains to be tested.

We also found an indirect way in which information in subspaces interfere with each other: divisive normalization of population activity. This led to a decrease of working memory information in Delay 2 once motor preparation information emerged. Divisive normalization, which has been described before in the LPFC^12,13^, could be useful as an energy saving mechanism, since it maintains the population activity at a low level when new information is added. A bump attractor model with divisive normalization allowed us to replicate the properties of LPFC activity. However, this model only provides a high-level support for this mechanism, in need of a mechanistic implementation.

In this work, we derived two subspaces, and analyzed the benefits of decoding from those subspaces, from data in which the memory location and the motor preparation location were identical. However, there are situations where the LPFC is required to store multiple pieces of information that are uncorrelated, e.g. if the animal has to remember the location and color of a target to perform a task, where these are uncorrelated^21,22^. We have verified that our approach can be extended to identify the relevant subspaces in tasks with uncorrelated information as well (see Methods). We show that in tasks with uncorrelated information, decoding in the full space could result in higher interference as compared to tasks with correlated information (Supplementary 15). However, the use of subspaces could reduce interference in both cases (Supplementary 15), suggesting a possible advantage of encoding information in independent subspaces in a broad range of cognitive tasks that we typically associate with the LPFC.

Our results support a framework in which low-dimensional subspaces could be a general property of cortical networks^6,23^. Under this framework, downstream regions could extract specific information from these subspaces^8^. This could provide a mechanism for selective routing of information between regions^24^, which could in turn be a building block of our cognitive flexibility capacity. Furthermore, this framework also provides intuitions underlying cognitive capacity limitations. The dimensionality of a network’s full state space constrains the number and dimensionality of the different information subspaces that could coexist within the network. The LPFC is a densely connected brain hub, anatomically connected to more than 80 regions, compared to the roughly 30 connected to the primary visual cortex^25^. Given the variety of inputs to the LPFC, it is not surprising that a large number of its neurons show mixed selective responses^1,2^, which in turn endow the LPFC with high dimensionality^1^. This property would allow a higher number of subspaces to coexist within the network.

## Acknowledgements

This work was supported by startup grants from the Ministry of Education Tier 1 Academic Research Fund and SINAPSE to C.L., a grant from the NUS-NUHS Memory Networks Program to S.-C.Y., and a grant from the Ministry of Education Tier 2 Academic Research Fund to C.L. and S.-C.Y. (MOE2016-T2-2-117).

## Author Contributions

S.-C.Y., C.T., and C.L., conceptualized the analysis framework. C.T., R.H., and A.P. performed the analysis. S.-C.Y. and C.L. guided the data analysis. All authors discussed the results, and C.T., S.-C.Y., and C.L. wrote the manuscript.

## Online Methods

### Subjects and Surgical Procedures

We used two male adult macaques (*Macaca fascicularis*), Animal A (age 4) and Animal B (age 6), in the experiments. All animal procedures were approved by, and conducted in compliance with the standards of the Agri-Food and Veterinary Authority of Singapore and the Singapore Health Services Institutional Animal Care and Use Committee (SingHealth IACUC #2012/SHS/757). The procedures also conformed to the recommendations described in Guidelines for the Care and Use of Mammals in Neuroscience and Behavioral Research (National Academies Press, 2003). Each animal was implanted first with a titanium head-post (Crist Instruments, MD, USA) before arrays of intracortical microelectrodes (MicroProbes, MD, USA) were implanted in multiple regions of the left frontal cortex. In Animal A, we implanted 6 arrays of 16 electrodes and 1 array of 32 electrodes in the LPFC, and 2 arrays of 32 electrodes in the FEF, for a total of 192 electrodes. In Animal B, we implanted 1 array of 16 electrodes and 2 arrays of 32 electrodes in the LPFC, and 2 arrays of 16 electrodes in the FEF, for a total of 112 electrodes. The arrays consisted of platinum-iridium wires with either 200 or 400 µm separation, 1 – 5.5 mm of length, 0.5 MΩ of impedance, and arranged in 4×4 or 8×4 grids. Surgical procedures followed the following steps. 24 hours prior to the surgery, the animals received a dose of Dexamethasone to control inflammation during and after the surgery. They also received antibiotics (amoxicillin 7 – 15 mg/kg and Enrofloxacin 5 mg/kg) for 8 days, starting 24 hours before the surgery. During surgery, the scalp was incised, and the muscles retracted to expose the skull. A craniotomy was performed (~ 2×2 cm). The dura mater was cut and removed from the craniotomy site. Arrays of electrodes were slowly lowered into the brain using a stereotaxic manipulator. Once all the arrays were secured in place, the arrays’ connectors were secured on top of the skull using bone cement. A head-holder was also secured using bone cement. The piece of bone removed during the craniotomy was repositioned to its original location and secured in place using metal plates. The skin was sutured on top of the craniotomy site, and stitched in place, avoiding any tension to ensure good healing of the wound. All surgeries were conducted using aseptic techniques under general anesthesia (isofluorane 1 – 1.5% for maintenance). The depth of anesthesia was assessed by monitoring the heart rate and movement of the animal, and the level of anesthesia was adjusted as necessary. Analgesics were provided during post-surgical recovery, including a Fentanyl patch (12.5 mg/2.5 kg 24 hours prior to surgery, and removed 48 hours after surgery), and Meloxicam (0.2 – 0.3 mg/kg after the removal of the Fentanyl patch). Animals were not euthanized at the end of the study.

### Recording Techniques

Neural signals were initially acquired using a 128-channel and a 256-channel Plexon OmniPlex system (Plexon Inc., TX, USA) with a sampling rate of 40 kHz. The wide-band signals were band-pass filtered between 300 to 3000 Hz. Following that, spikes were detected using an automated Hidden Markov Model based algorithm for each channel^26^. The eye positions were obtained using an infrared-based eye-tracking device from SR Research Ltd. (Eyelink 1000 Plus). The behavioral task was designed on a standalone PC (stimulus PC) using the Psychophysics Toolbox^27^ in MATLAB (Mathworks, MA, USA). In order to align the neural and behavioral activity (trial epochs and eye data) for data analysis, we generated strobe words denoting trial epochs and performance (rewarded or failure) during the trial. These strobe words were generated on the stimulus PC, and were sent to the Plexon and Eyelink computers using the parallel port.

### Behavioral Task

Each trial started with a mandatory period (500 ms) where the animal fixated on a white circle at the center of the screen. While continuing to fixate, the animal was presented with a target (a red square) for 300 ms at any one of eight locations in a 3×3 grid. The center square of the 3×3 grid contained the fixation spot and was not used. The presentation of the target was followed by a delay of 1000 ms, during which the animal was expected to maintain fixation on the white circle at the center. At the end of this delay, a distractor (a green square) was presented for 300 ms at any one of the seven locations (other than where the target was presented). This was again followed by a delay of 1,000 ms. The animal was then given a cue (the disappearance of the fixation spot) at the end of the second delay to make a saccade towards the target location that was presented earlier in the trial. Saccades to the target location within a latency of 150 ms and continued fixation at the saccade location for 200 ms was considered a correct trial. An illustration of the task is shown in Figure 1a. One of the animals was presented with only 7 of the 8 target locations because of a behavior bias in the animal.

### Cross-Temporal Decoding

In order to assess the stability of the population code, we used data at each time point to train a decoder based on Linear Discriminant Analysis (LDA), built using the classify function in MATLAB, and tested the decoder on data from other time points as described in Parthasarathy et al. 2017^2^. When decoding data in the full space, we denoised the training and testing data using principal components analysis (PCA) at every time point by reconstructing the data with the top n principal components that explained at least 95% of the variance. In order to decode data in the subspace, the PCA projection matrix described in the previous step was replaced by the matrix specifying the desired subspace (working memory or motor preparation), and the resulting data in the subspace would thus have 7 dimensions.

### Pattern decorrelation and subspace identification

In this analysis, we used a pseudo-population of *N = 226* neurons. For each trial *condition* (which was one of seven possible target locations), we trial-averaged and time-averaged the neural activity in Delay 1 (800 to 1,300 ms from target onset) and Delay 2 (2,000 to 2,500 ms from target onset) to obtain two activity matrices of *226 x 7*. We then normalized the two activity matrices to the mean of the baseline by subtracting neural responses in the fixation period (300 ms before target onset), and obtained activity matrices 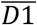 and 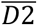 of size *226 x 7*, where each column represented the change in population activity under one condition. 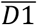 and 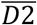 contained activity with high correlation (Supplementary Fig. 1a,b). We looked for the decorrelated components 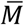 and 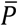 (Supplementary Fig. 1c,d), which were also of size *226 x 7*, such that 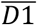 and 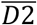 were the linear combination of 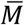 and 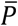 with different mixing coefficients, which could be formulated as:

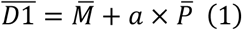

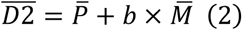

where *a* and *b* were scalars. We can rewrite 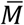 and 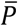 as:

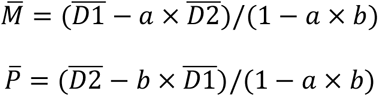

and each pair of (*a*, *b*) will determine one pair of 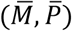. We concatenated 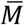 and 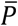, respectively, across conditions, and evaluated the independence between 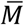 and 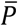 by computing the mutual information between the two arrays of size 1582. We performed a parameter search for (*a*, *b*) in the range of [−2, 2] with a step size of 0.01. The minimum mutual information obtained was 0.076 bits when *a* = 0.12 and *b* = 0.65 (Supplementary Fig. 1). The memory and preparation subspaces were then defined by the bases of 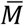 and 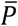.

A similar decorrelation was performed between activity in Delay 1 and the pre-saccadic period (150 to 0 ms prior to saccade onset):

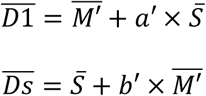

where 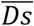 was the activity in the pre-saccade period. Using the same approach as described above, we obtained 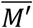 and 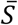 with a minimum mutual information of 0.086 when *a*’ = 0.01 and *b*’ = 0.71 (Supplementary Fig. 6).

### Angle between subspaces

The subspaces we were comparing had dimensions of *226 x 7* (Supplementary Fig. 5). In order to calculate the angle between two subspaces, we first computed the inner product matrix of size *7 x 7*, took the mean of the absolute values to get a scalar value, and lastly, converted this scalar value to angle by using the inverse cosine function. In 226 dimensions, the angles between 2 random vectors were almost all compressed in the range from 80 to 90 degrees, so the angle by itself did not provide a good indication of similarity between two subspaces with high dimensions. In order to get an unbiased estimation of the similarity of two subspaces in high dimensions (i.e. *Mem* and *Stable* in Suppl. Fig. 5), we first computed the angle between *Mem* and *Stable*, and then computed the angle between *Stable* and a random subspace of the same dimension, and lastly, computed the difference between the two angles. By repeating this process, we generated a distribution of the pairwise difference between the two angles. If the 95th percentile of this distribution was smaller than 0, we considered the angle between *Mem* and *Stable* to be significantly smaller than chance.

### Neural Network decoding

In theory, the maximum decodable information in the full state space cannot be lower than the decodable information in a lower-dimensional subspace. However, using LDA, we found that the decoding accuracy in the full space was lower compared to that in the working memory and motor preparation subspaces (compare Fig. 1b and d,e). The reason for this was that LDA overfitted the training set due to noise and sparse samples in the full space, which led to lower performance on the test set. In order to more reliably compare the decodable information in the full space and the subspaces for working memory and motor preparation, we used a neural network classifier that can reduce the issue of overfitting by introducing a validation set and early stopping in the training process. The data were split into non-overlapping training (50%), validation (30%), and testing (20%) sets. In each iteration of the training, we trained the decoder on the training set, and evaluated it on the validation set. Once the validation result began to decrease (indicating overfitting on the training set), we stopped the training, and reported the decoder’s performance on the test set. The model had a simple architecture, with only one hidden layer of 10 units (with a linear transfer function), and one output layer (Softmax) for pattern recognition.

In order to extract the working memory and motor preparation activity in the full space with single-trial variability, we used a variation of Equation (1) and (2):

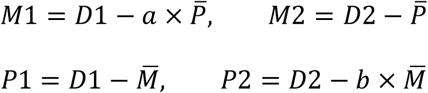

where 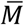 and 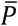 are the trial-averaged memory and preparation component, *D*1 and *D*2 were the single-trial Delay 1 and Delay 2 activity matrices of size *226 x 1750* (250 random single trials per condition); *M* and *P* (number indicates in Delay 1 and Delay 2) were the single trial working memory and motor preparation activity in the full space (of size *226 x 1750*). Subspace memory and prepara̅tion we̅re obtained by projecting the full space activity into their respective subspaces defined by 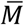 and 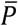.

In the error trial analysis, error trial full space memory and preparation activity were estimated by:

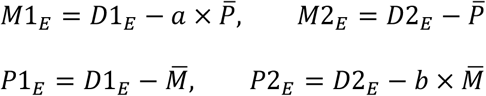

where *D*1_*E*_ and *D*2_*E*_ were similar to *D*1 and *D*2, but from error trials. The decoder was trained and validated on the data from correct trials in the subspace and tested on the data from error trials in the same subspace. Although we interpreted the decrease in decoding performance in the two subspaces in error trials to be evidence of the link between these subspaces and the behavior of the animal, an alternative interpretation could be that there was a general increase in noise in the population in error trials (perhaps due to factors like inattention), and this led to a non-specific decrease in information in all subspaces, including the memory and preparation subspaces. However, due to the lack of target information in the null space, this non-specific decrease in information in the null space would not be observable because of the floor effect. In order to rule this possibility out, we quantified the intra-cluster variance in the full space across locations for correct and error trials in both Delays 1 and 2 (refer to Supplementary Figure 7). We found no evidence supporting the fact that the intra-cluster variance in Delay 1 was higher in error trials than in correct trials (P > 0.46, g = 0.85), and found the intra-cluster variance in error trials in Delay 2 to actually be lower than correct trials (P < 0.01, g = 6.6), presumably due to the effects of divisive normalization. These results indicated that the drop in performance in the working memory and motor preparation subspaces in error trials was not due to a non-specific increase in noise, but were more likely due to the fact that the responses in error trials deviated significantly from those in correct trials, resulting in lower information in the two subspaces.

### Identification of null space

The memory and preparation subspaces both had dimension of *226 x 7* (i.e. 7 vectors in a 226 dimensional space). We used MATLAB function *null()* to find the complementary space of the conjunctive subspace for both memory and preparation. The null spaces had rank of 212, and were orthogonal to both working memory and motor preparation subspaces. 117 PCs in the null space could explain 95% of the total variance after projection the full space activity into the null space. Any information about the target location that was not captured by the two subspaces would be captured by the null space.

### Statistics

We considered two bootstrapped distributions to be significantly different if the 95th percentile range of the two distributions did not overlap. We also computed an estimated p-value for this comparison using the following formula^28^,

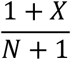

where *X* represents the number of overlapping data points between the two distributions and *N* used throughout the paper, two distributions with no overlap will result in a p-value < 0.001, and two distributions with x% of overlap will result in a p-value ~ x/100.

In addition to the estimated p-value, we also computed the effect size of the comparison using a measure known as Hedges’ g, computed using the following formula^29^,

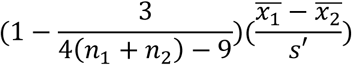

where

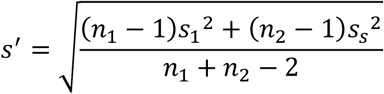

*X* refers to the mean of each distribution, n refers to the length of each distribution, and s refers to the standard deviation of each distribution.

No statistical methods were used to pre-determine sample sizes, but our sample sizes were similar to those reported in previous publications^16,30,31^. The majority of our analyses made use of nonparametric permutation tests, and as such, did not make assumptions regarding the distribution of the data. No randomization was used during the data collection, except in the selection of the target and distractor locations for each trial. Randomization was used extensively in the data analyzed to test for statistical significance. Data collection and analysis were not performed blind to the conditions of the experiments. No animals or data points were excluded from any of the analyses. Please see additional information in the Life Sciences Reporting Summary.

### Cell selectivity classification

For Supplementary Figure 10, and in order to match the selectivity properties of neurons in the model with those of LPFC data, we first quantified the selectivity of LPFC activity as follows. First, using a two-way ANOVA with independent variables of target locations (7 locations) and task epoch (Delay 1 and Delay 2), we categorized cells as those with *pure working memory* selectivity (those with target information in both Delay 1 and Delay 2, and those with selectivity to target location and task epoch, but no interaction, 27.6% of cells), those with *mixed* selectivity to target location and task epoch (those with a significant main effect of target location and task epoch, as well as a significant interaction between target location and task epoch, 43.9% of cells). And using two one-way ANOVAs of target location (one in Delay 1 and one in Delay 2), we categorized cells as those with *pure motor preparation* selectivity (those with significant selectivity in Delay 2, but not Delay 1, 28.6% of cells).

### Model

For the bump attractor model, we used two populations of firing-rate units for the memory and preparation input (N = 80 for each), and tested the model’s performance with different overlapping ratios between the two populations (if the overlapping ratio was 0%, then the full network consisted of 160 units, whereas if the overlapping ratio was 100%, then the full network consisted of 80 units). We used simplified discrete time equations to describe the dynamics of the population:

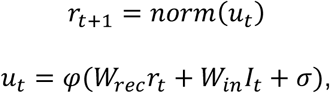

where *r*_*t*_ was the population firing rate at time *t*, *W*_*rec*_ was the recurrent connection weight between units, *I*_*t*_ was the external input at time *t*, *W*_*in*_ was the loading weight of input signal to the population, *σ* ∼ *N*(0,0.1) was a noise term, *norm*(*x*) was a divisive normalization function that kept the mean population firing rate at the baseline level, and *φ*(*x*) was a piecewise nonlinear activation function adopted from Wimmer et al., 2014^15^:

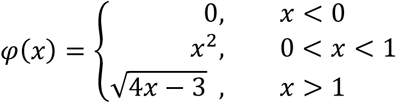

The matrix, *W*_*rec*_, had a diagonal shape with stronger positive values near the diagonal, and weaker negative values elsewhere, such that only a few neighboring units were connected via excitatory weights to each other while being connected via inhibitory weights to the rest. In this way, a structured input signal to adjacent units was able to generate a local self-sustaining bump of activity. There were eight input units, representing the eight spatial target locations in the animal’s task. For each input unit, the loading weight matrix *W*_*in*_ specified a random group of 10 adjacent units in the working memory population as well as the motor preparation population to receive the signal. The input to the working memory population was transiently active in the target presentation period, and the input to the motor preparation population was transiently active in the distractor presentation period. The whole population consisted of the working memory and motor preparation populations, and normalization was applied on the whole population.

For the linear subspace model, we used a total of N = 112 units where the dynamics of the activity could be described as:

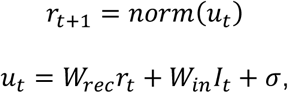

where *r*_*t*_ was the population firing rate at time *t*: *W*_*rec*_ was the recurrent connection weight between units; *I*_*t*_ was the external input at time *t*; *W*_*in*_ was the loading weight of the input signal including 0 or 0.1, denoted as N (0,0.1)); and *norm*(*x*) was a divisive normalization function that kept the mean population firing rate at the baseline level. We constructed the recurrent weight matrix from eigendecomposition:

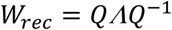

where *Q* was a random square matrix whose columns were the eigenvectors of *W*_*rec*_, and *Λ* was a diagonal matrix whose diagonal elements were the corresponding eigenvalues for each eigenvector. The first 17 eigenvalues in *Λ* were 1 (thus there were 17 stable eigenvectors), while the rest of the eigenvalues were randomly chosen between 0 and 1 using a uniform distribution. In each simulation, we assigned 1 stable eigenvector for baseline activity (with entries selected from a uniform distribution U(0,1)), 8 stable eigenvectors for working memory activity, and 8 stable eigenvectors for motor preparation activity (with entries selected from U(1,2)). In order to ensure that decoding performance in Delay 1 and Delay 2 were the same, we imposed a positive mean for the motor preparation activity, so that the incorporation of motor preparation in Delay 2 would elevate the population mean, and divisive normalization would reduce the mean activity of both working memory and motor preparation information. Otherwise, if the motor preparation activity had zero mean, there would be a significant increase of decoding performance in Delay 2. In the input weight matrix, the input activity for working memory corresponded to the 8 working memory eigenvectors, and the input activity for motor preparation corresponded to the 8 motor preparation eigenvectors. The distractor inputs had the same input activity as did the target inputs, but with a lower magnitude (0.2 compared to target). At the beginning of each trial, the population started with baseline activity equal to the stable baseline eigenvector, then the input for working memory was transiently active in the target presentation period, and the input for motor preparation was transiently active in the distractor presentation period. As all the input activity corresponded to stable eigenvectors, all target information, distractor information, and motor preparation information were maintained stably across time.

#### Subspaces for uncorrelated information

Due to our experimental design, the working memory location and the motor preparation locations were identical in each trial, and thus correlated. We can imagine another case where there are two types of information that do not have a one-to-one mapping (for example, in a task that requires memorizing locations of items - 1 out of 2 possible locations, and their colors - 1 out of 3 possible colors, which are uncorrelated). When each target location is associated with only one stimulus color (similar to our working memory and motor preparation locations), the incorporation of stimulus color information in Delay 2 would add only 1 out of 3 possible shifts (representing the 3 possible stimulus colors) to the clusters representing target location (Supplementary Fig. 15a). However, when target location and stimulus color are uncorrelated (each stimulus color is equally likely to appear in each target location), the incorporation of stimulus color information could add any of the 3 possible shifts to the clusters representing target location activity, leading to much more diffuse clusters (Supplementary Fig. 15b). In this latter case, we propose a more general formulation to estimate the information subspaces for target location and stimulus color. First, we group̅̅t̅rials by target location and obtain the trial-averaged and time-averaged activity in Delay 1 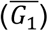. Next, we group trials by stimulus color and obtain the trial-averaged and time-averaged activity in Delay 2 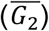. Finally we estimate the subspaces by:

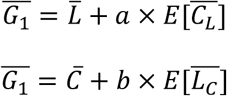

where 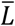 and 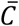 define the subspaces for target location and stimulus color, while *a and b* are scalars representing the mixing coefficients. 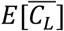 represents the expectation of stimulus color activity associated with particular target locations, and 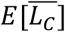 represents the expectation of target location activity associated with particular stimulus colors. At one extreme, the correlation between target location and stimulus color could be 0 (completely random pairing), in which case the expectation value will reduce to 0 if averaging across the other variable does not result in a net translation (which also means there will be no code morphing). In this case, there is no need to minimize mutual information, as the 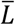 and 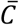 vectors will remain unchanged. On the other hand, if there is a net translation, code morphing will be present, and there will be a need to minimize the mutual information to recover the angles between the subspaces. At the other extreme, the correlation could be 1 (one-to-one mapping), in which case the expectation value will reduce to 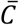 and 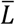. We can verify that the decorrelation method used for working memory and motor preparation components in this work was a special case of this formulation (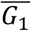 and 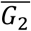 become 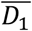 and 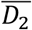 as memory and preparation have the same grouping, and the expectations reduce to 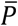 and 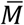 as there is one-to-one mapping). We would perform the same parameter search on (*a, b*) that will give the least mutual information between 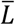 and 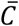.

## Data Availability

The data that support the findings of this study are available from the corresponding authors upon reasonable request.

## Code Availability

A code package for analyses used in the paper is available at: https://github.com/chengtang827/MemoryPreparationSubspace

---

**Supplementary Figure 1.**
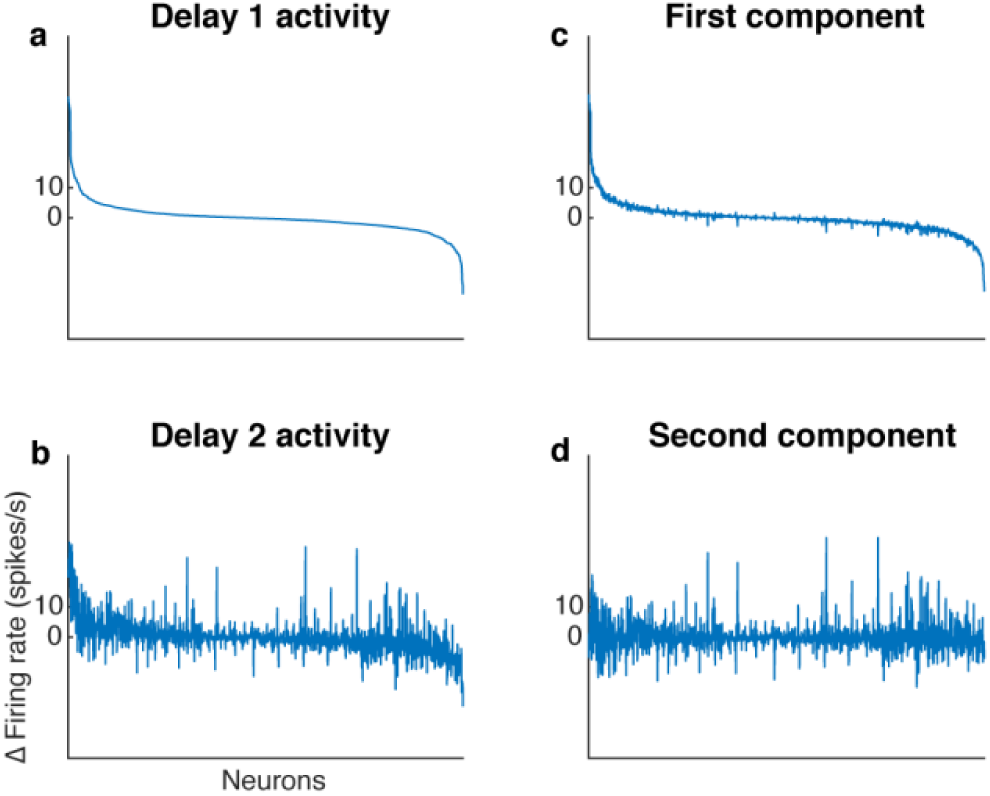
Decorrelated population activity between Delay 1 and Delay 2. **a**, Delay 1 population activity for all seven target locations sorted according to firing rate. The x-axis has 1582 points (226 cells x 7 locations). Each neuron’s firing rate for the last 500 ms of Delay 1 on each trial was averaged across time, before being averaged across trials in each location. This was then subtracted by the average baseline firing rate (averaged across 300 ms prior to target presentation before averaging across trials). The neurons were sorted in descending order by the Delay 1 activity in all four plots (a to d). **b**, Delay 2 population activity, significantly correlated with Delay 1 activity (*r* = 0.69, *P* < 0.001, mutual information = 0.33 bits). **c**, The decorrelated *first component*. **d**, The decorrelated *second component*, has minimal mutual information with the *first component* (*r* = 0.017, *P* = 0.5, mutual information = 0.076 bits).

**Supplementary Figure 2.**
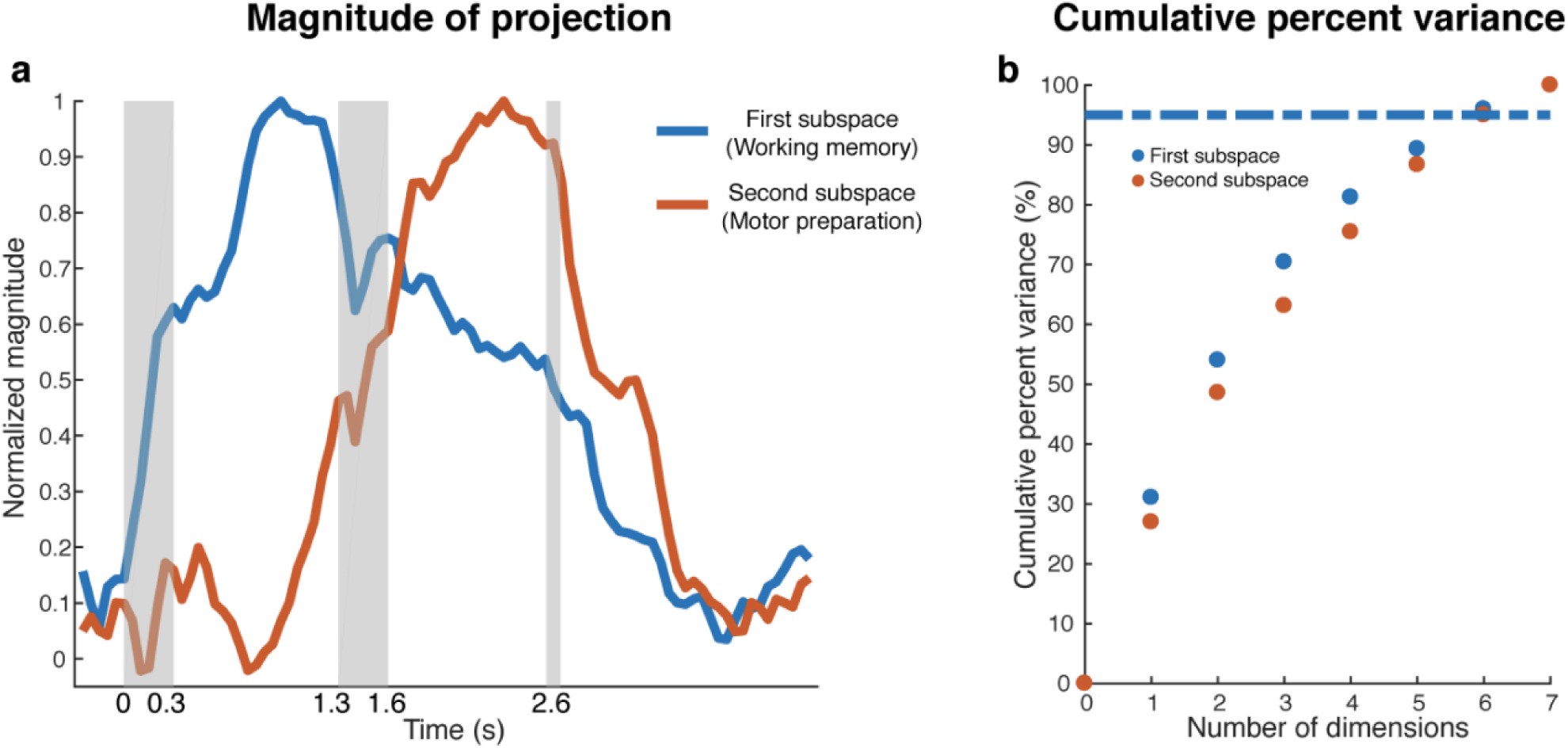
Magnitude of projections in continuous time bins and effective dimension. a, We projected the trial-averaged full space population activity for each time bin across the whole trial into the first subspace and the second subspace, and calculated the magnitude of the projections. For each subspace, the magnitude was normalized to have a maximum value of 1. The projections into the first and second subspaces exhibited different temporal profiles. b, We projected the full space activity from both Delay 1 and Delay 2 (time-averaged in each period) into the two subspaces, and calculated the cumulative percent variance explained by the principal components in each projection. In both subspaces, 6 PCs were needed to explain more than 95% of the variance.

**Supplementary Figure 3.**
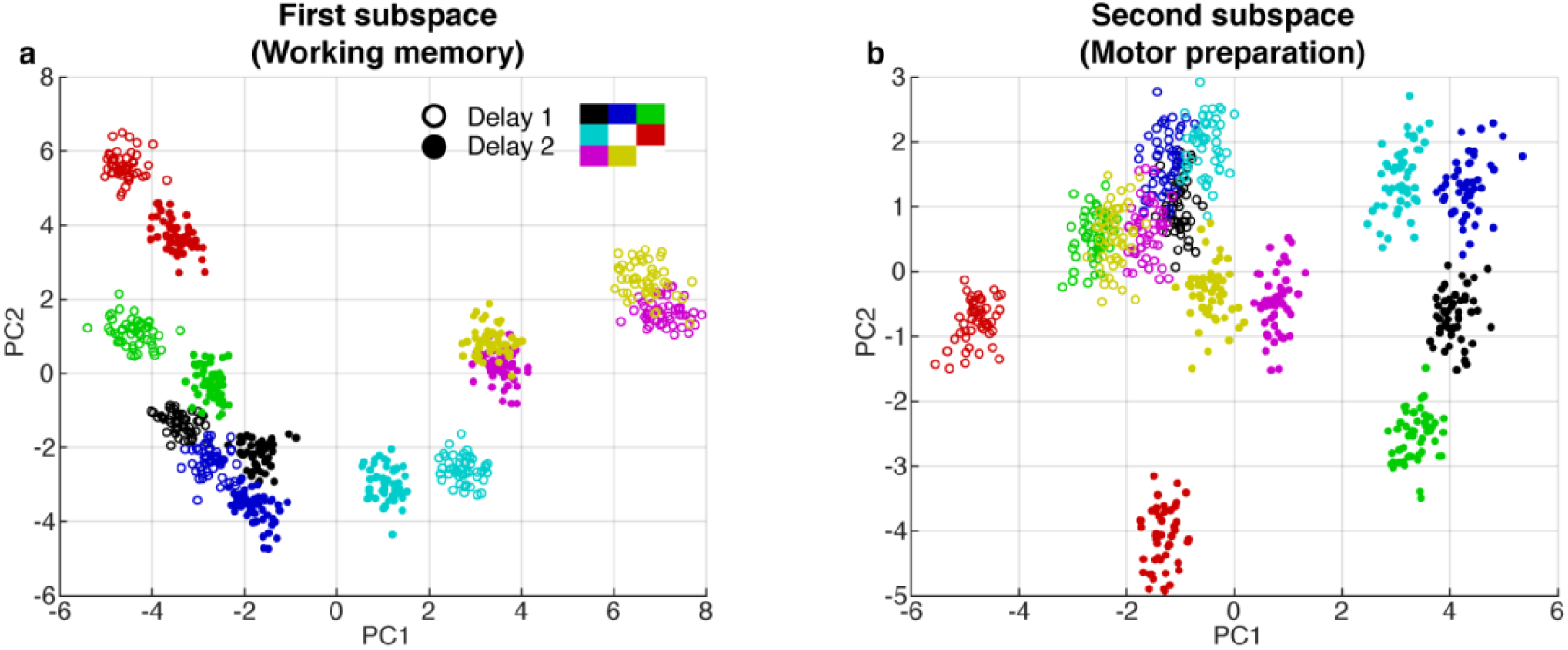
PCA projections in the first and second subspaces. **a**, Delay 1 and Delay 2 activity for 50 trials at each target location projected into the first subspace (top 2 PCs). Open circle, Delay 1 activity; closed circle, Delay 2 activity. Target locations are color coded according to the legend. Delay 2 clusters appeared to move closer to each other compared to the Delay 1 clusters. This meant that the boundaries of the classifiers trained in Delay 2 would work better for Delay 1 activity, compared to the opposite scenario, as seen in the off-diagonal quadrants in Figure 1d. **b**, Delay 1 and Delay 2 activity projected into the motor preparation subspace (top 2 PCs). The clusters exhibited significant overlap during Delay 1, but separated into distinct clusters in Delay 2.

**Supplementary Figure 4.**
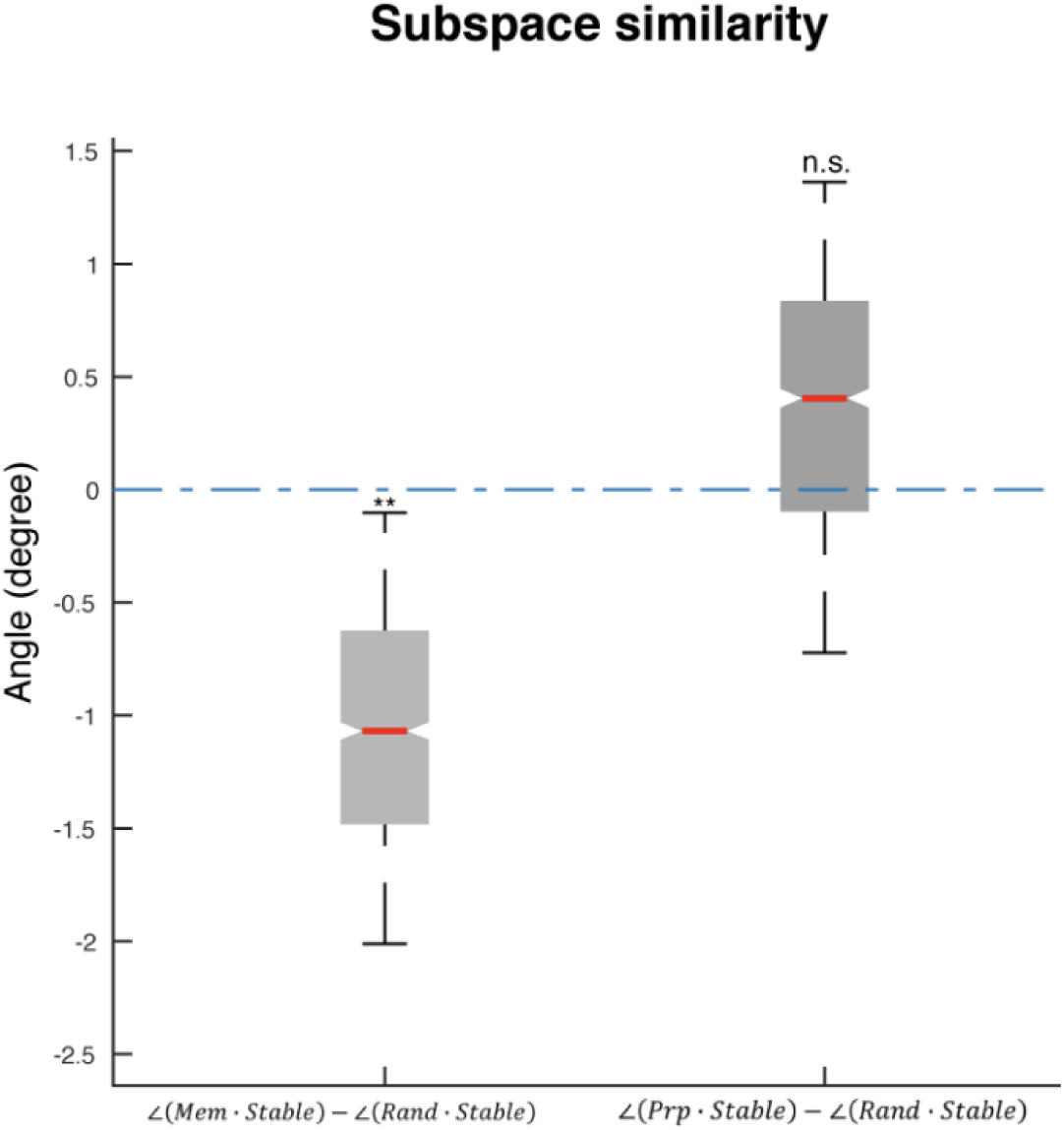
Angle between subspaces. The following abbreviations are used in this figure: Mem - the working memory subspace identified in this work; Prp - the motor preparation subspace identified in this work; Stable - the stable memory subspace identified in Parthasarathy et al. 2019 using an optimization method. Due to bootstrapping of pseudo-trials, different runs of the optimization gave slightly different results; and Rand - a random subspace generated from a 226 x 65 PCA space that explained 95 percent of the variance in the full space delay activity. Each column (226 x 1) of the random subspace (226 x 7) was a random linear combination of the 65 PCs, and each column was normalized to unit length. Left boxplot, For each of the 1000 Stable subspaces we generated, we computed the pairwise difference between its angle with the working memory subspace and a random subspace. The working memory subspace was significantly closer to the stable subspace than chance, providing support that the memory subspace was similar to the stable memory subspace. Right boxplot, For each Stable subspace, we computed the pairwise difference between its angle with the motor preparation subspace and a random subspace. The motor preparation subspace was not significantly closer to the stable subspace than chance, providing support that the motor preparation subspace was unrelated to the stable subspace. Asterisks (**), significant (i.e., 95th percentile range of the distribution did not overlap with zero); n.s., nonsignificant (i.e., 95th percentile range of the distribution overlapped with zero).

**Supplementary Figure 5.**
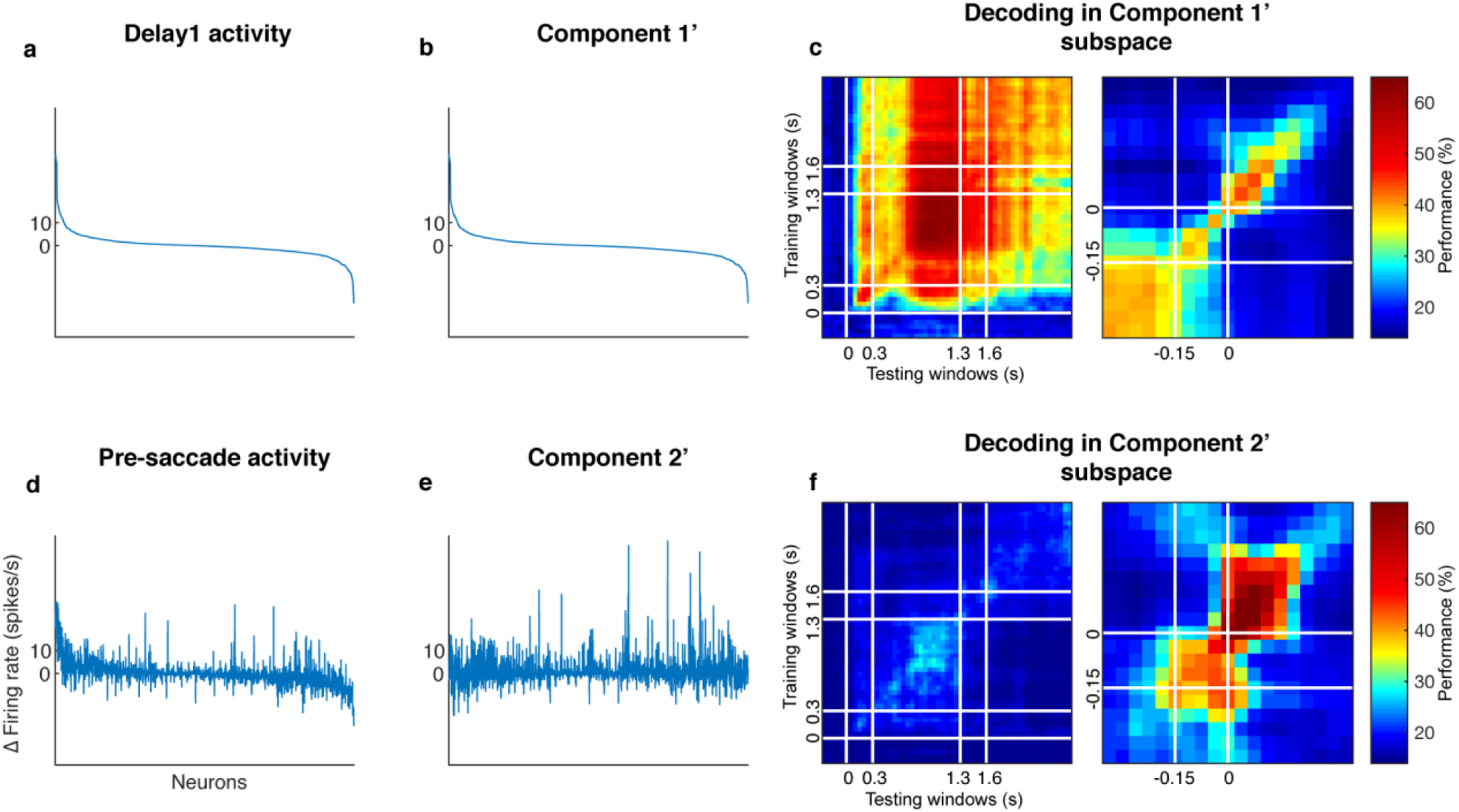
Decorrelated population activity between Delay 1 and the pre-saccadic period. **a**, Delay 1 population activity concatenated for all seven target locations. The x-axis has 1582 points (226 cells x 7 locations). Each neuron’s firing rate was time-averaged and trial-averaged for each of the 7 locations, and subtracted by its baseline firing rate. The index was sorted in descending order by the Delay 1 activity in Panels a, b, d, and e. **b**, The decorrelated Component 1’. **c**, Cross-temporal decoding after projecting the full space activity into the subspace defined by Component 1’. Left panel, aligned to target onset. Right panel, aligned to saccade onset. The white lines indicate the 150 ms window used to obtain the pre-saccade response used in the decorrelation. **d**, Pre-saccade period activity, significantly correlated with Delay 1 activity (*r* = 0.50, *P* < 0.01, mutual information = 0.19 bits). **e**, The decorrelated Component 2’, has minimal mutual information with Component 1’ (*r* = −0.018, *P* = 0.49, mutual information = 0.087 bits). **f**, Cross-temporal decoding after projecting the full space activity into the subspace defined by Component 2’. Left panel, aligned to target onset, right panel, aligned to saccade onset.

**Supplementary Figure 6.**
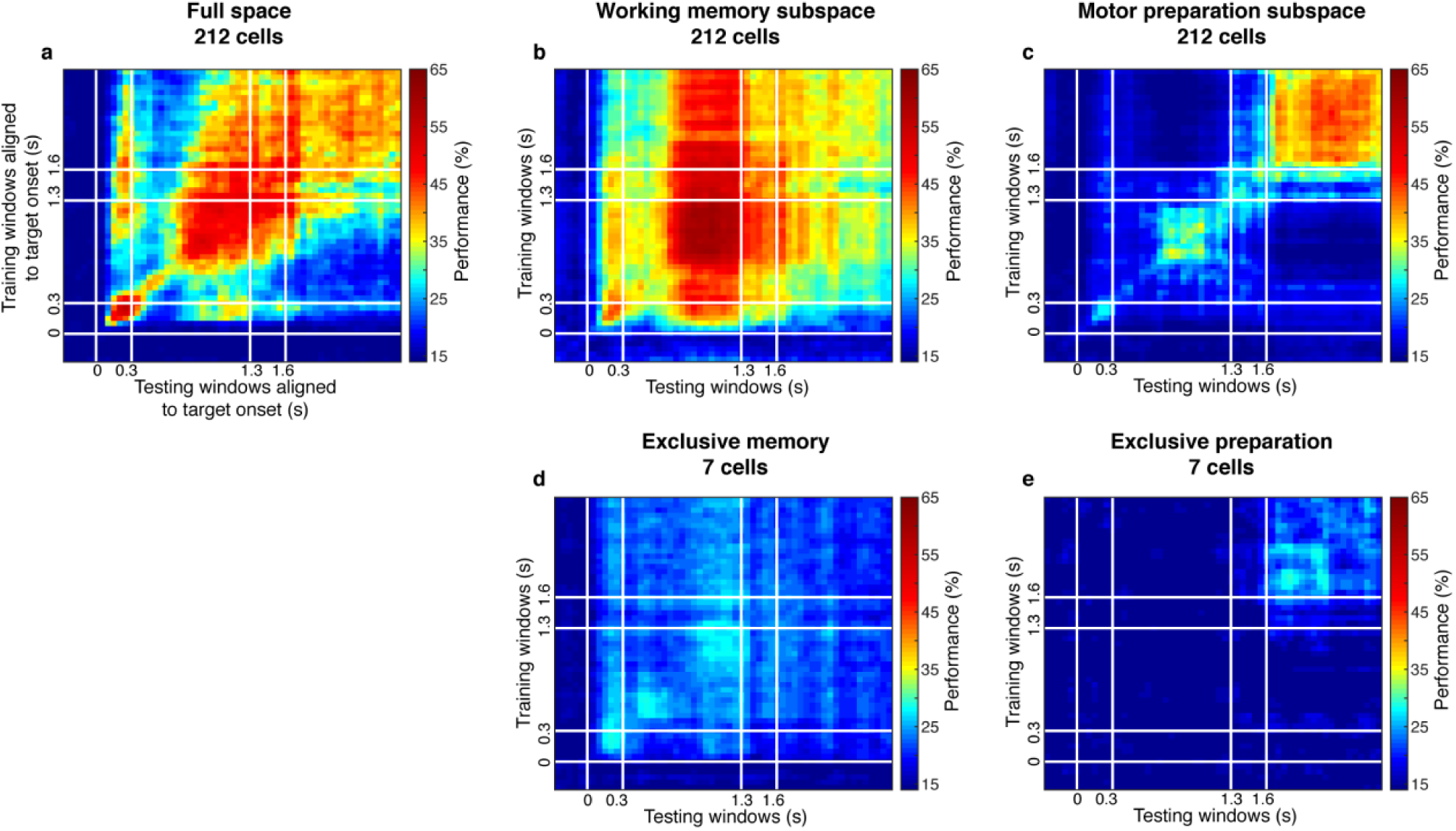
Cross-temporal decoding on the population with mixed selectivity and populations with exclusive selectivity. **a**, Cross-temporal decoding of the 212 cells (within the 2 standard deviations in the ratio distribution shown in Fig. 3b) in the full space. **b**, Cross-temporal decoding of the 212 cells in the working memory subspace identified on the 212 cells using the same decorrelation method. **c**, Cross-temporal decoding of the 212 cells in the motor preparation subspace identified on the 212 cells using the same decorrelation method. **d**, Cross-temporal decoding of the 7 cells with exclusive loading into the working memory subspace. **e**, Cross-temporal decoding of the 7 cells with exclusive loading into the motor preparation subspace.

**Supplementary Figure 7.**
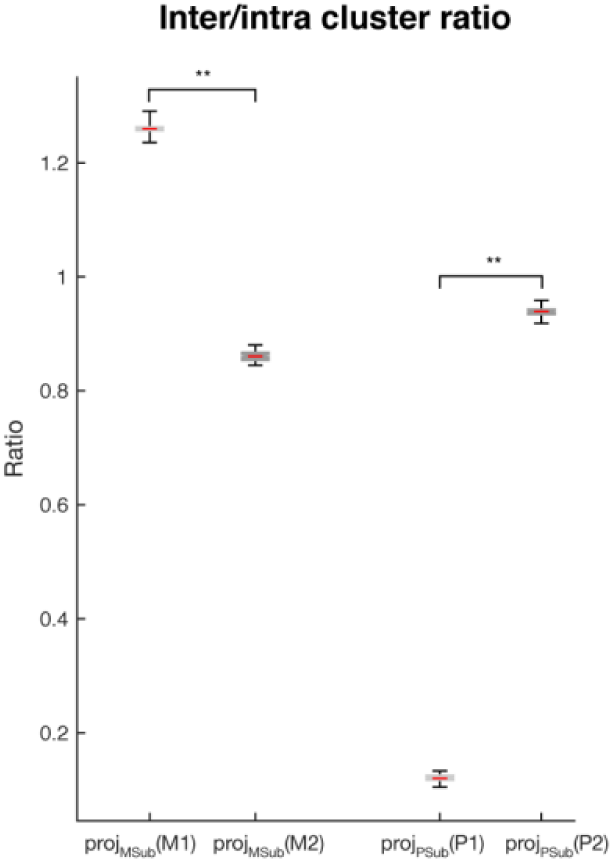
Inter- and intra-cluster distance analysis. The ratios of the inter-cluster distance (the average of the pairwise Euclidean distance between cluster means) and the intra-cluster distance (the average of the pairwise distances between samples within each cluster, which were then further averaged across clusters) are shown for: *proj*_*MSub*_*(M1)* - projection of the working memory component in Delay 1 into the working memory subspace; *proj*_*MSub*_*(M2)* - working memory component in Delay 2 projected into the working memory subspace; *proj*_*PSub*_*(P1)* - motor preparation component in Delay 1 projected into the motor preparation subspace; and *proj*_*PSub*_*(P2)* - motor preparation component in Delay 2 projected into the motor preparation subspace. Asterisks (**), significant (i.e., 95th percentile range of the two distributions did not overlap).

**Supplementary Figure 8.**
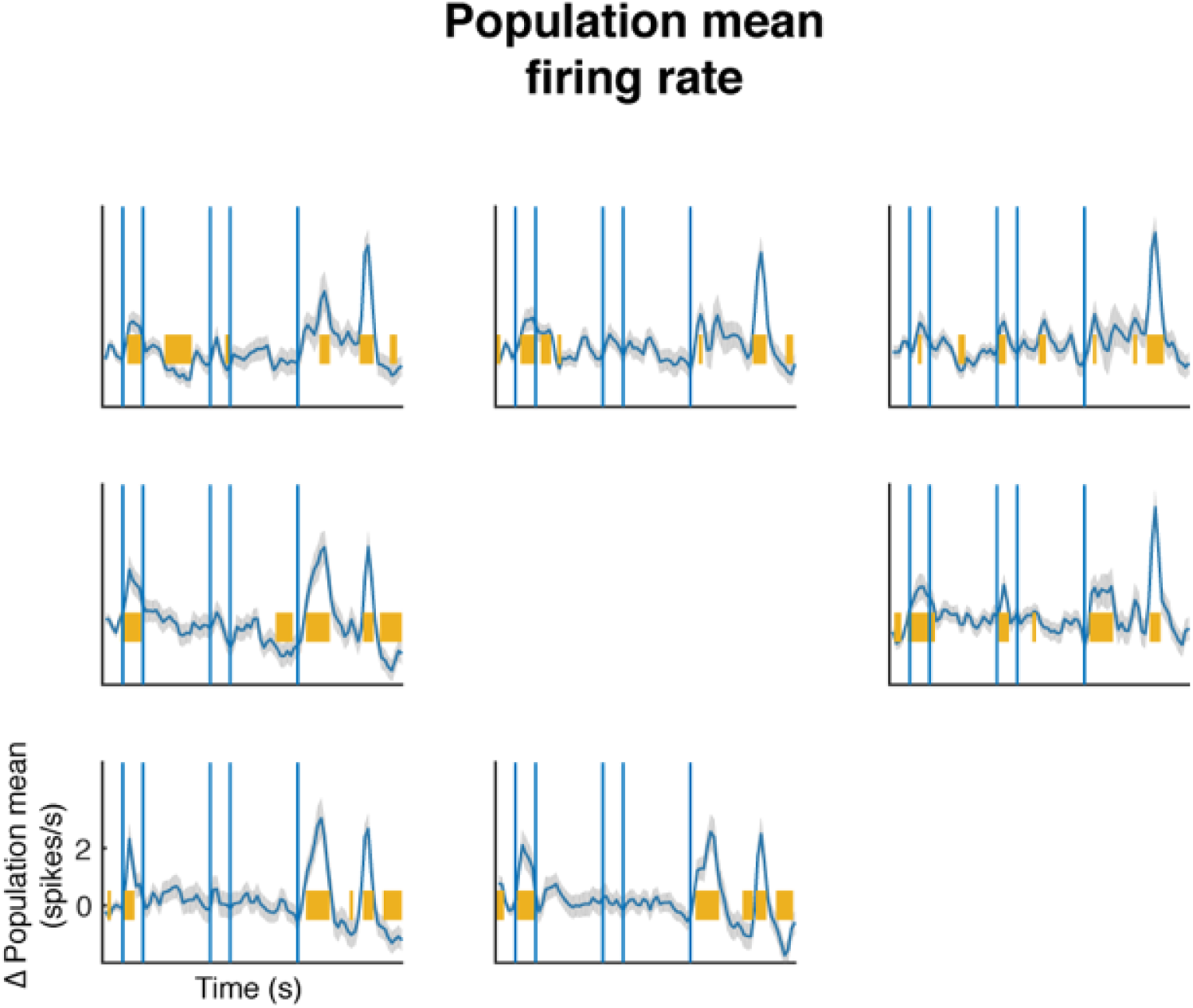
Mean population firing rate. We trial-averaged each cell’s firing rate in each target condition in each time bin, and subtracted each cell’s baseline firing rate (mean of the fixation period, which was 300 ms prior to target presentation). The blue line indicates the mean firing rate of the population (226 cells). The shaded area represents the standard error. The yellow line indicates time bins in which the population firing rate was significantly different from zero (T-test, *P* < 0.05).

**Supplementary Figure 9.**
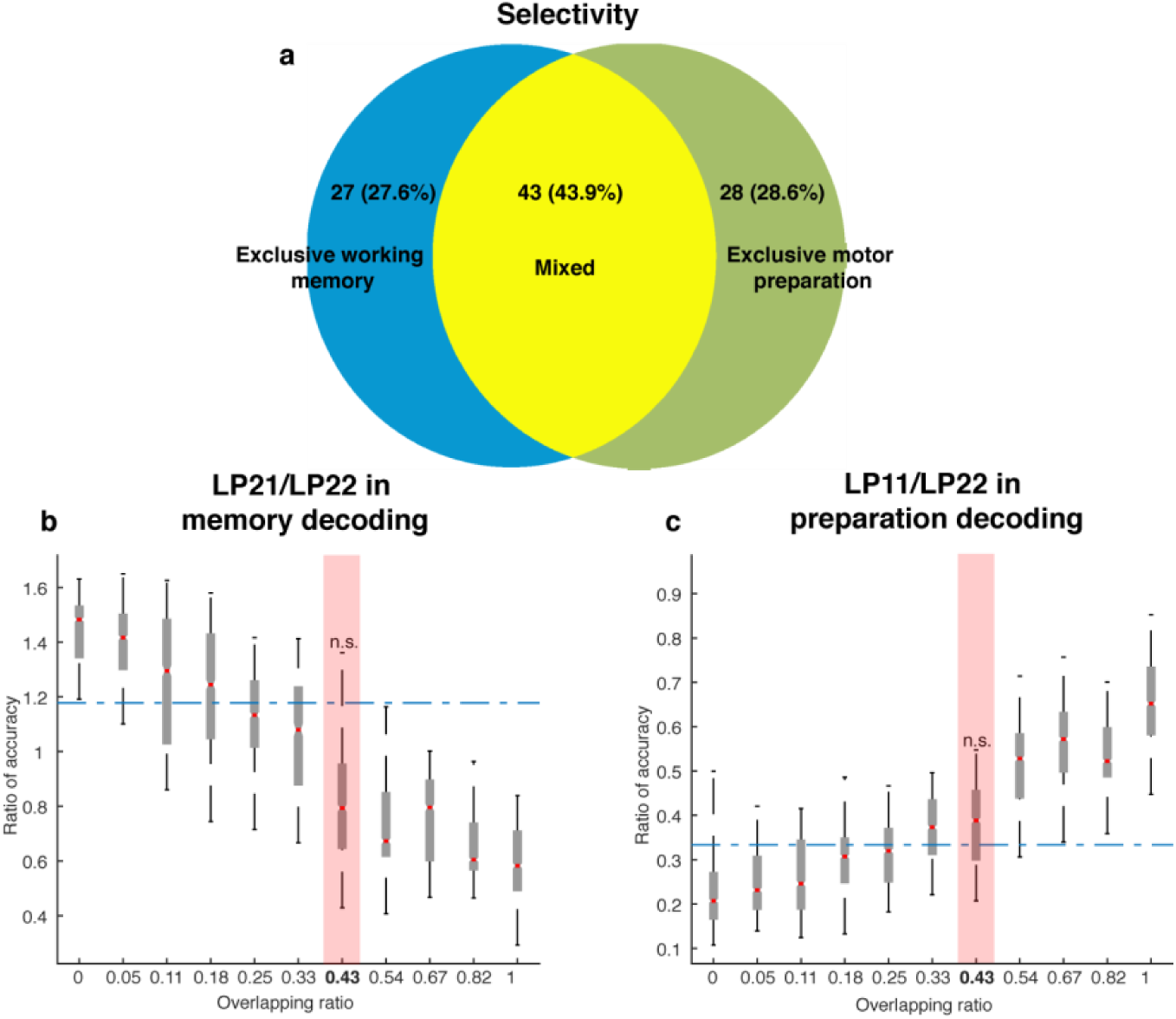
Neuronal selectivity. **a**, In order to classify the selectivity of individual neurons, we used a two-way ANOVA with independent variables of target locations (7 locations) and task epoch (Delay 1 and Delay 2) to categorize cells as: 1) those with *exclusive working memory* selectivity (those with target information in both Delay 1 and Delay 2, and with selectivity to target location and task epoch, but no interaction, 27.6% of cells); and 2) those with *mixed* selectivity to target location and task epoch (those with a significant main effect of target location and task epoch, as well as a significant interaction between target location and task epoch, 43.9% of cells). Additionally, we used two one-way ANOVAs of target location (one in Delay 1, and one in Delay 2) to categorize cells as those with *exclusive motor preparation* selectivity (those with significant selectivity in Delay 2, but not Delay 1, 28.6% of cells). Among the cells that exhibited selectivity in the delay periods (98 cells), we estimated that 27.6% had *exclusive working memory* selectivity, 28.6% had *exclusive motor preparation* selectivity, and 43.9% had *mixed* selectivity to both working memory and motor preparation. **b**, Comparison of the decoding in the working memory subspace between neural data (blue dashed line) and model data (box plots). X-axis: different population overlapping ratios between working memory and motor preparation, where overlapping ratio was the number of cells with *mixed* selectivity divided by the total population size. Neural data showed an overlapping ratio of *0.43*. Y-axis: mean decoding accuracy in LP21 divided by mean accuracy in LP22 (refer to Fig. 5e). **c**, Comparison of the decoding in the preparation subspace between neural data (blue dashed line) and model data (box plots). X-axis, same as in Panel a. Y-axis, mean decoding accuracy in LP11 divided by mean accuracy in LP22 (refer to Fig. 5f). n.s., nonsignificant (i.e., 95th percentile range of the distribution overlapped with the dashed line representing the result from the neural data).

**Supplementary Figure 10.**
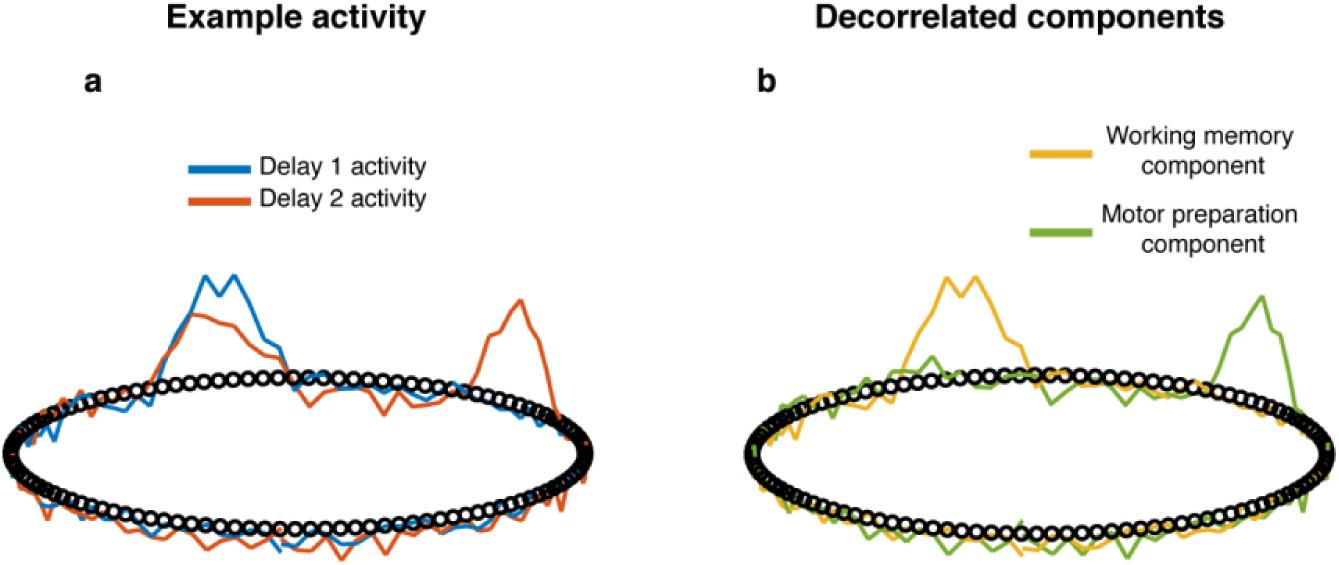
Model with normalization **a**, An example of the activity found in the model units in one trial. There was one bump in Delay 1 (blue trace), and two bumps in Delay 2 (red trace). Note that the overlapping bump in Delay 2 was smaller, which was a result of divisive normalization. **b**, The memory (yellow trace) and preparation (green trace) components demixed by decorrelating Delay 1 and Delay 2 activity.

**Supplementary Figure 11.**
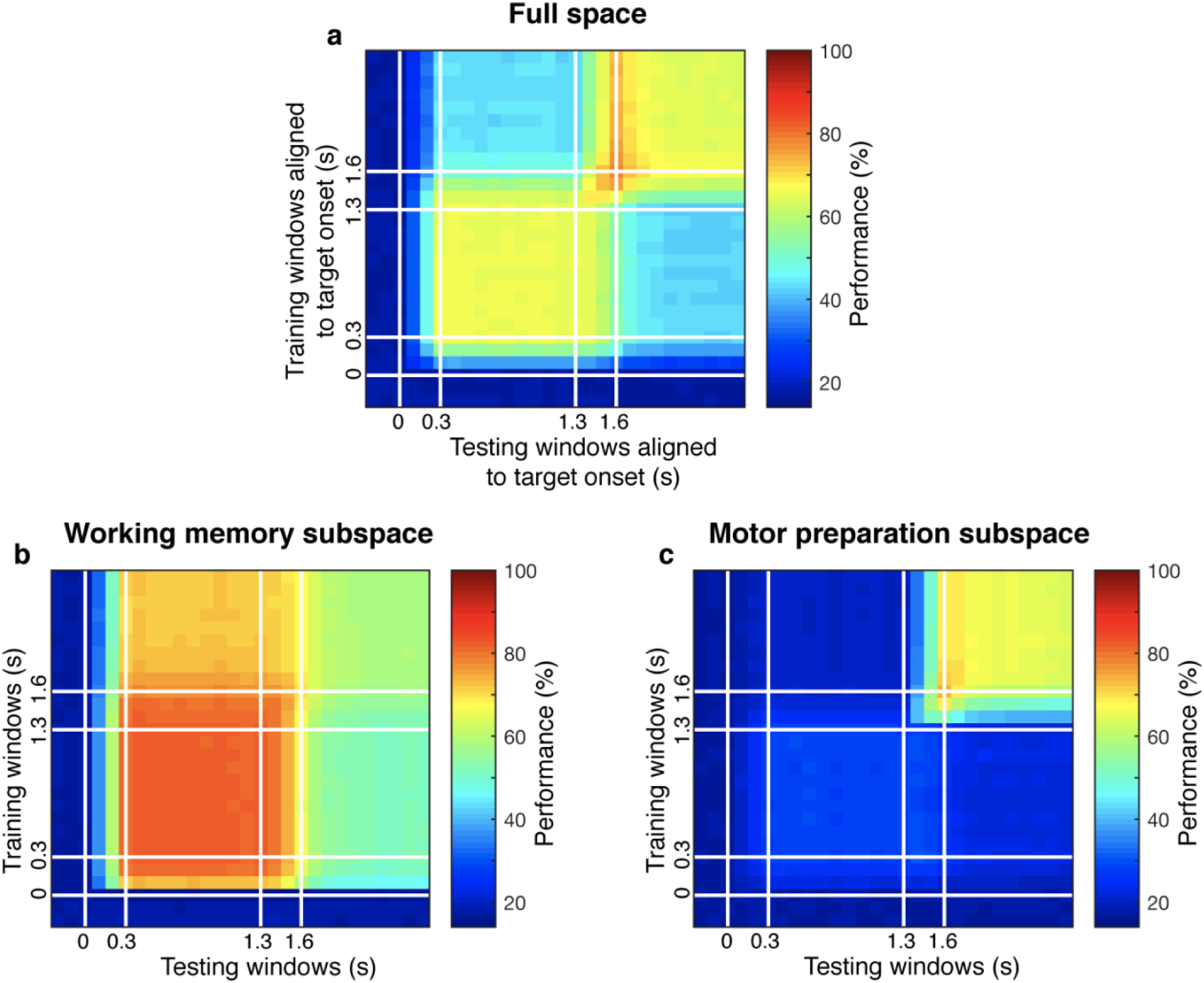
Linear subspace model a, In the linear subspace model, working memory information was encoded by 8 stable eigenvectors, and motor preparation information was also encoded by 8 stable eigenvectors (see Methods). For each target location, there was a one-to-one mapping of working memory activity and motor preparation activity. Code morphing was replicated in the full space. b, The decay of working memory information was replicated in the working memory subspace identified by the decorrelation method. c, The emergence of motor preparation information was replicated in the motor preparation subspace identified by the decorrelation method. We believe that the bump attractor model and linear subspace model share some conceptual similarities in our case: both models have mechanism to maintain stable activity in the absence of sustained external input, and are able to incorporate new information without affecting existing information. Further, bump attractor model could be deemed as a special case of linear subspace model where the reciprocally excitatory units are grouped together, which has more biologically support.

**Supplementary Figure 12.**
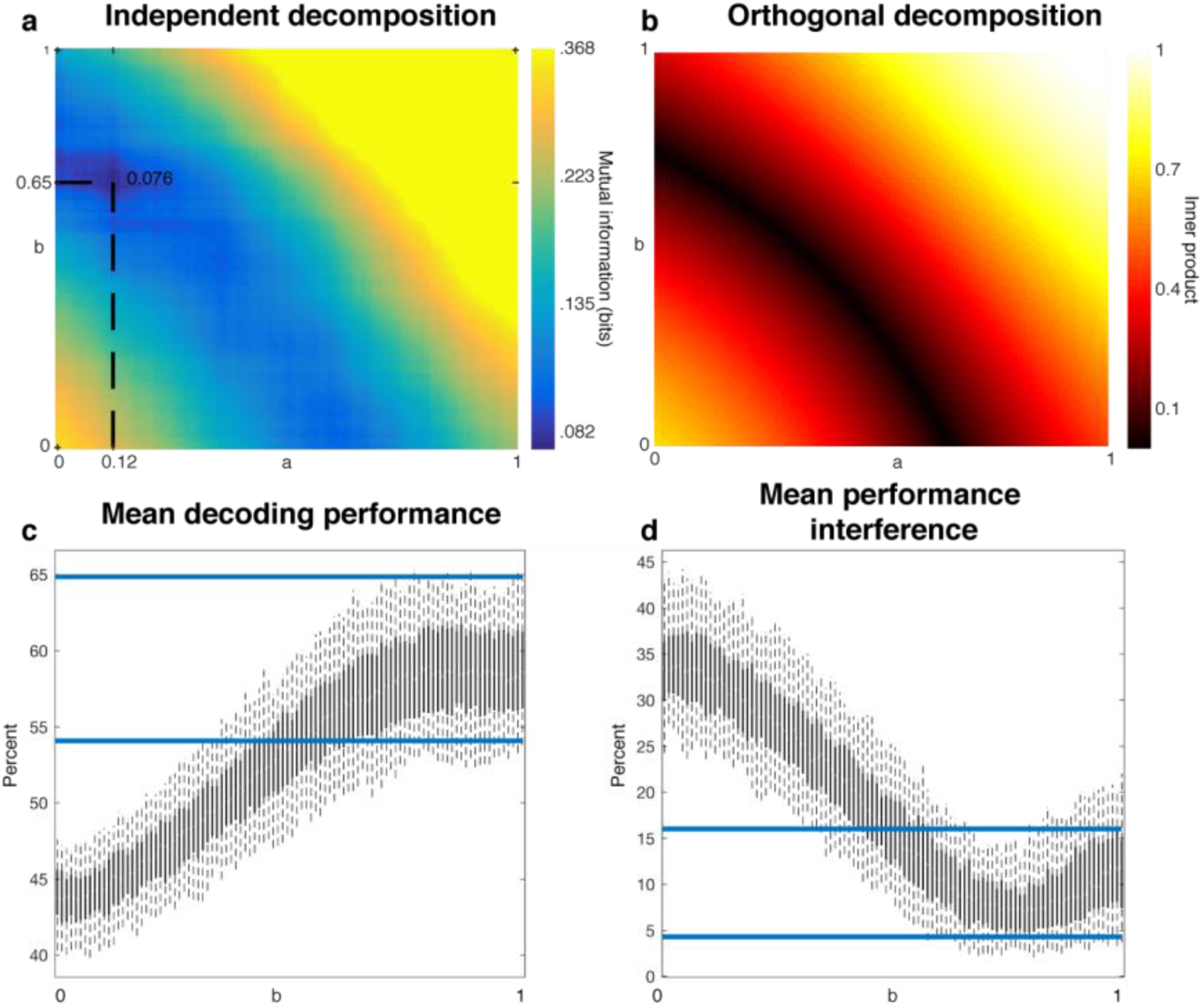
Independent and orthogonal decomposition. **a**, Heatmap showing the mutual information between working memory and motor preparation components with different mixing coefficients. There was a global minimum when a = 0.12 and b = 0.65. **b**, Heatmap showing the inner product between working memory and motor preparation components with different mixing coefficients. There was no unique global minimum. **c**, The mean decoding performance, defined as (proj_MSub_(M) + proj_PSub_(P))*/2* in Delay 2 (refer to Fig. 4), is shown for different values of b from Panel b that combined with their respective values of a to return the minimum inner product. The range of the values of b was normalized to [0,1]). We bootstrapped half of the total population 1000 times, and calculated the results for each bootstrap. Each bar was the 5^th^ and 95^th^ percentile of the bootstrap results for a particular value of b. The blue lines are the 5^th^ and 95^th^ percentile of the decoding performance achieved using the global minimum of the mutual information decomposition shown in Panel a. The orthogonal decomposition did not provide significantly better performance than the independent decomposition. **d**, Mean performance interference was defined as ((proj_MSub_(M) – proj_MSub_(M+P)) + (proj_PSub_(P) *–* proj_PSub_(M+P)))/2 in Delay 2 (refer to Fig. 4a,b). The rest are the same as in Panel c.

**Supplementary Figure 13.**
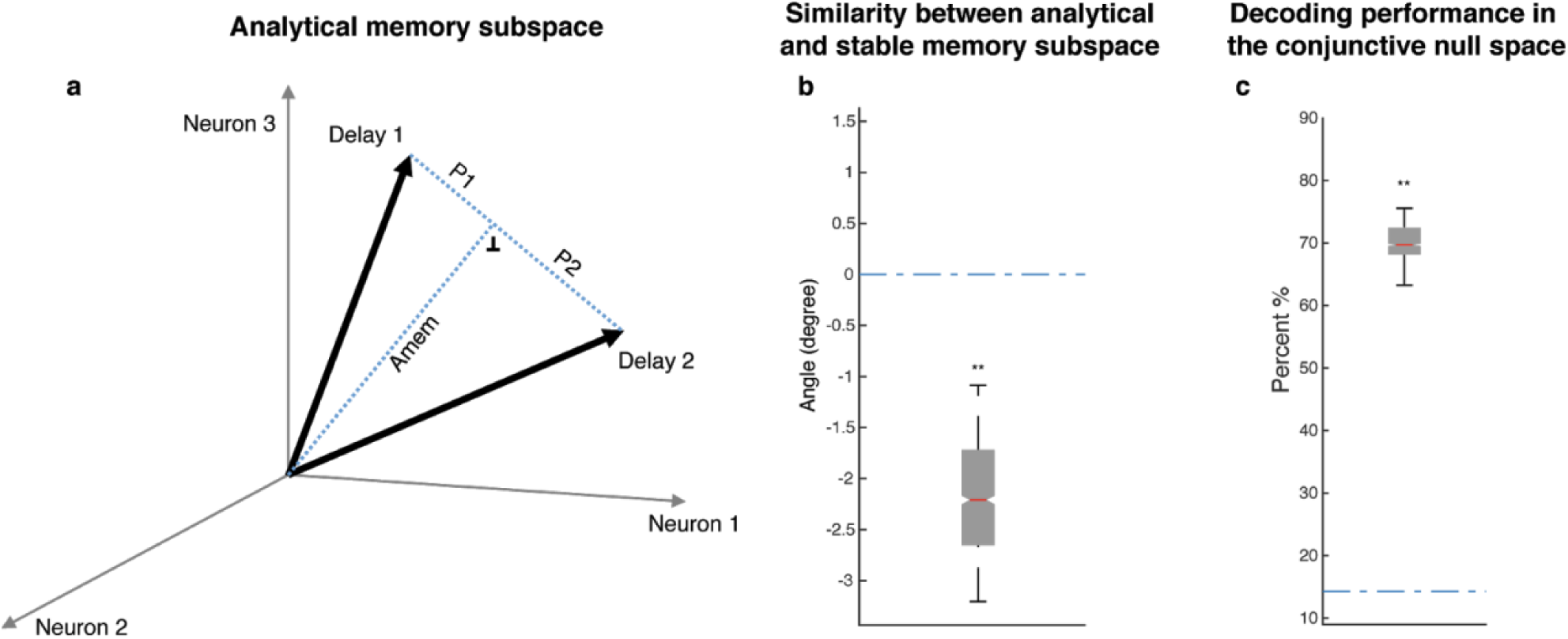
Analytical memory subspace and non-memory subspace. a, We calculated the difference vector between Delay 2 and Delay 1 activity (Delay 2 - Delay 1), and defined the null space of the difference vector as a stable memory subspace (which we called the Analytical memory subspace, Amem), such that the projection of Delay 1 and Delay 2 activity into this subspace overlapped. We calculated the preparation component as the residual vector between D1, D2, and Amem, i.e. D1 - Amem, and D2 - Amem, respectively. However, this would result in a preparation component with the same coefficients but have opposite signs in Delay 1 and Delay 2 (P1 and P2 in the figure). In other words, that implied that an ‘anti-preparation’ signal to the target location had to exist in Delay 1, and then was inverted to the “true preparation” signal after distractor presentation. This seemed unnecessarily complicated, and required the existence of an “anti-signal” before the “signal” even emerged, which seemed unlikely for a cognitive process. b, In order to investigate the similarity between Amem and the stable memory subspace identified in Parthasarathy et al. 2019 using an optimization method, we generated 1000 stable memory subspaces, and computed the difference in angle between each of the stable memory subspaces with Amem, and the difference in angle between the stable memory subspace and a random subspace (refer to Supplementary Figure 5). The stable memory subspace was significantly closer to Amem than chance, providing support that the stable memory subspace was similar to Amem. **c**, In Parthasarathy et al. 2019, we postulated a ‘non-memory’ input that was the same for all target locations that, together with a stable memory subspace, was able to capture the code morphing and the prevalence of neurons with non-linear mixed selectivity. Here, we defined the non-memory subspace as the mean of Delay 2 - Delay 1 vectors across all target locations, and performed a similar analysis as in Figure 4c in the conjunctive null space of the working memory and non-memory subspaces. Unlike the working memory and motor preparation subspaces, the working memory and non-memory subspaces were not able to capture all the target information in Delay 2 (there was significant information in the conjunctive null space), indicating that the motor preparation subspace was a better fit to the neural data than the ‘non-memory’ input.

**Supplementary Figure 14.**
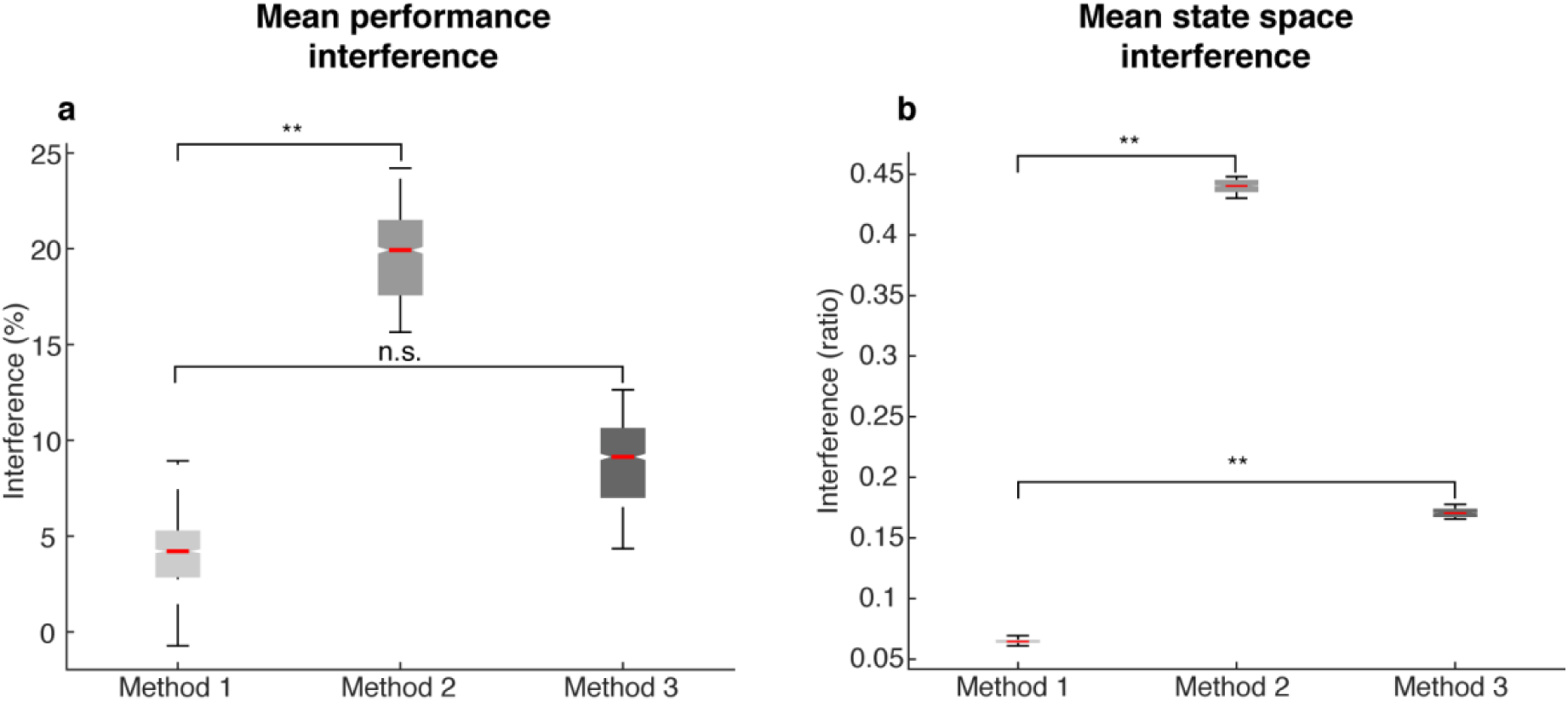
Amount of interference in different methods. The following labels are used in this figure: ***Method 1*** -by minimizing mutual information, we decorrelated population activity in Delay 1 and Delay 2, and found working memory and motor preparation components with the least mutual information; ***Method 2*** -defined Delay 1 activity as the memory component, and the subtraction of Delay 1 from Delay 2 (D2 - D1) as the preparation component. This method resulted in two components that were highly correlated, and thus showed a larger interference between subspaces compared to the interference found using Method 1; and ***Method 3*** -the Analytical memory subspace defined in Supplementary Figure 13. **a**, Mean performance interference, as defined as in Supplementary Figure 12. Method 1 showed significantly lower levels of mean performance interference than Method 2 (*P* < 0.001, *g* = 5.87), but was not significantly different from Method 3 (*P* > 0.65, *g* = 1.86). **b**, Mean state space interference, defined as the inter/intra-cluster ratio for ((proj_MSub_(M) – proj_MSub_(M+P)) + (proj_PSub_(P) *–* proj_PSub_(M+P)))/2 in Delay 2 (refer to Fig. 4d,e). Both Method 2 and Method 3 showed significantly higher levels of state space interference than Method 1. Asterisks (**), significant (i.e., 95th percentile range of the two distributions did not overlap). n.s., nonsignificant (i.e., 95th percentile range of the two distributions overlapped).

**Supplementary Figure 15.**
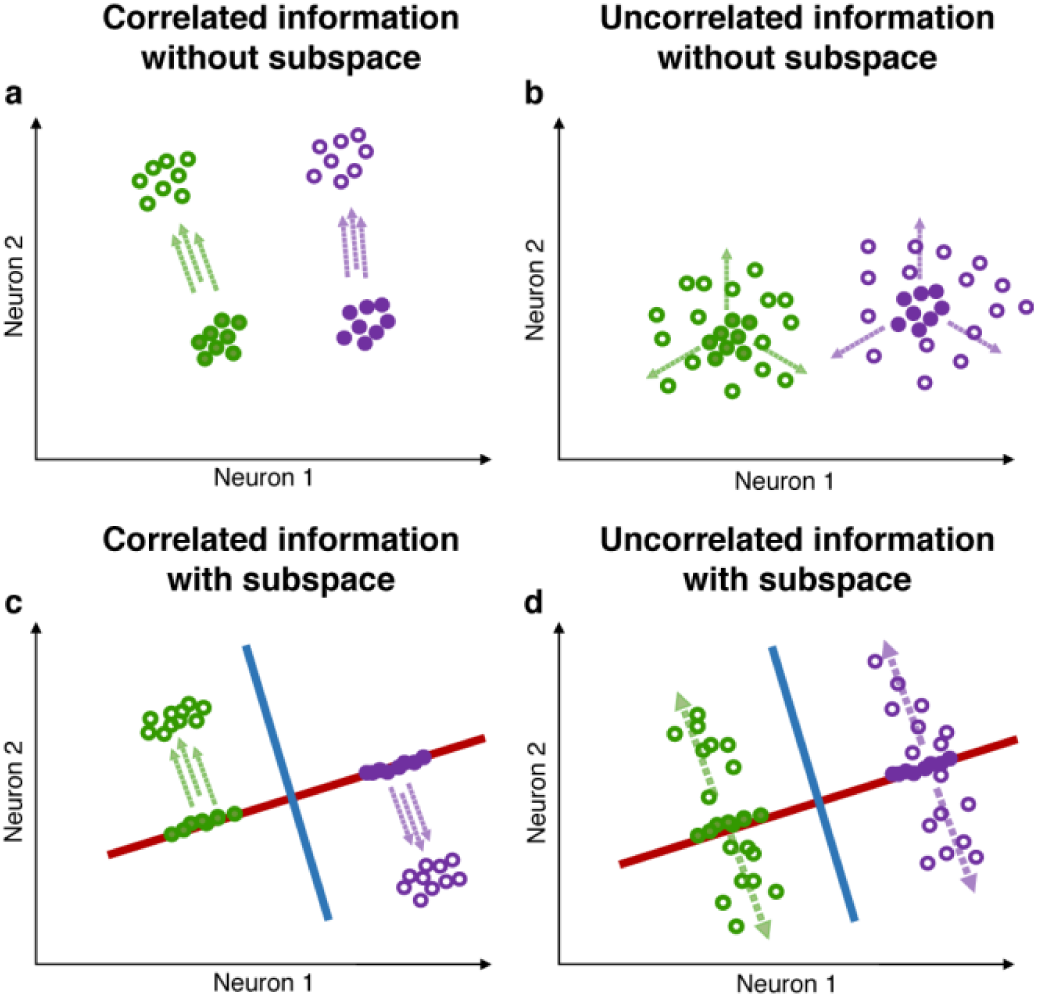
Correlated and uncorrelated information. **a**, Illustration of correlated information (in this example, 2 possible target locations and 3 possible stimulus colors, one target location is associated with only one stimulus color). Green and purple circles represent neuronal activity grouped by different target locations. Closed circles represent trial epochs containing only location information, while open circles represent trial epochs with color information incorporated into the location information. When trials are grouped by locations, the addition of correlated color activity will shift the location clusters only in the direction indicated by the parallel dashed arrows, and may not result in a decrease in the ratio of inter/intra-cluster distance. **b**, In the case where target location and stimulus color are uncorrelated (each stimulus color is equally likely to appear in each target location), the addition of uncorrelated color activity (indicated by the 3 dashed arrows) will ‘diffuse’ clusters representing target location, thus resulting in a decrease in the ratio of inter/intra-cluster distances. **c** and **d**, Red and blue lines represent the location and color subspaces. In both scenarios (correlated and uncorrelated information), independent location and color subspaces will alleviate the interference between the two pieces of information, as the ratio of inter/intra-cluster distance in one subspace is largely unaffected by changes in activity in the other subspace.

